# Maternal PFAS Transfer through Lactation: Dolphin Milk Reveals Routes of Early-Life Exposure

**DOI:** 10.1101/2025.05.02.651903

**Authors:** Kara M. Joseph, Ashlee T. Falls, James N. Dodds, Michael L. Power, Erin S. Baker

## Abstract

Per- and polyfluoroalkyl substances (PFAS) continue to increase in concentration and prevalence in the environment due to the creation of emerging PFAS and lack of breakdown of legacy compounds. PFAS are known to both bioaccumulate and biomagnify, therefore, species higher on the food chain, such as marine mammals, are highly exposed to these chemicals. Although studies suggest that considerable maternal transfer of persistent organic pollutants occurs via lactation, data is lacking on the temporal trends associated with PFAS exposure. Here, we utilized a set of dolphin breastmilk samples from an individual mother across a two-year lactation period to evaluate longitudinal trends in PFAS concentrations and profiles. Thirty-six PFAS were detected using a multidimensional platform combining liquid chromatography, ion mobility spectrometry, and mass spectrometry (LC-IMS-MS), and of these, 17 PFAS were detected continuously across the nursing window of 103-706 days. Quantitative analysis specifically showed concentrations of perfluorooctanesulfonic acid (PFOS) alone surpass weekly intake recommendations from the European Food Safety Authority and Food Standards Australia New Zealand by 1,000-fold but decreased slightly over time, possibly due to transfer from feedings. Non-targeted analysis also identified 13 additional compounds including two long-chained perfluorosulfonic acids not traditionally targeted, as well as the PFOS precursors, perfluoroethylcyclohexane sulfonate (PFECHS) and 2-(N-ethylperfluorooctanesulfonamido)ethyl phosphate (SAmPAP). This study therefore suggests that breastmilk is a major contributor to early-life PFAS exposure to mammals, particularly for long-chained PFAS.

**SYNOPSIS:** High concentrations of PFAS in dolphin milk reflect bioaccumulation and biomagnification, highlighting their persistence and harm to aquatic ecosystems and breastfed offspring.

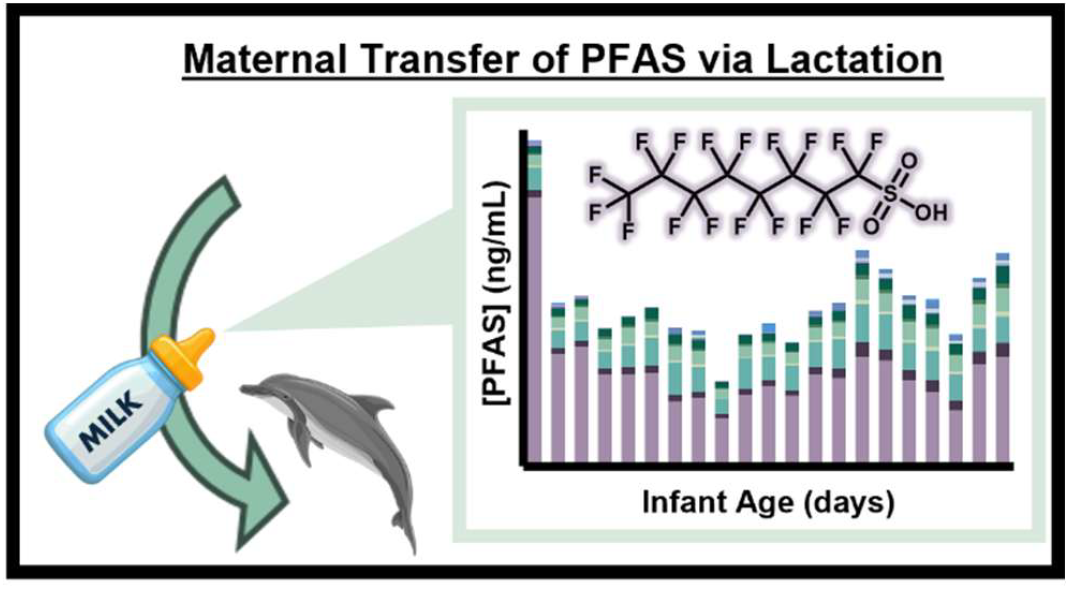

## 1. INTRODUCTION

Oceans are critical biomes, playing key roles in the maintenance of global biodiversity, nutrient cycling, and climate regulation. One threat to these habitats is a class of man-made chemicals called per- and polyfluoroalkyl substances (PFAS), which are widely used in a variety of consumer and industrial applications. PFAS belong to a class of compounds known as persistent organic pollutants (POPs), which encompass a range of environmentally ubiquitous chemicals that are toxic to humans and wildlife.^1, 2^ To date, more than 7 million compounds meet the current definition of a PFAS, any chemical containing a saturated CF_2_ or CF_3_ unit, with most having no toxicity information.^3, 4^ Due to their chemical composition, many PFAS are resistant to thermal and chemical degradation, bioaccumulate within individual organisms, and their concentrations are magnified at higher tropic levels.^1^ Since scientists have recently suggested that humanity has surpassed the planetary boundary for PFAS, major uncertainties must be addressed. However, current analytical capabilities struggle to keep up with the rapid production and spread of these chemicals, leaving significant gaps in our understanding of their distribution in ocean biomes and their effects on aquatic ecosystems. Therefore, we must continue to develop new ways to assess how PFAS interfere with marine organism health.

One common method to assess ecosystem health is the monitoring of one or more indicator species, which are animals that serve as proxies to the overall health of an ecosystem. Marine mammals are historically touted as ideal indicators of POP exposure in aquatic environments and can serve as early warning signs of ecosystem decline due to chemical pollution.^5-8^ Numerous studies have shown PFAS accumulate in the tissue and blood of marine mammals.^9-11^ Those most impacted are apex predators such as whales, seals, walruses, dolphins, and even Weddel seal populations in Antarctica, where PFAS concentrations frequently surpass what is considered safe for human exposure.^10, 12-17^ In humans, PFAS exposure is associated with increased risk of cancer, reduced immune response, lower birth weight, and dyslipidemia.^18-21^ In marine mammals, exposure to PFAS has been linked to similar effects such as perturbations in cell cycle, changes to liver and kidney function, and hematopoietic and immune system alterations.^22, 23^ Neonatal marine mammals are a particularly vulnerable population to these adverse effects, as it is widely accepted that breastfeeding results in a high concentration dose of PFAS during critical stages of growth and development for neonates, while causing a reduction in maternal PFAS body-burden.^24, 25^ Similar trends in high neonatal exposure to other POPs, such as polychlorinated biphenyls (PCBs), dichlorodiphenyldichloroethylene (DDE), and hexachlorobenzene (HCB) have been demonstrated via paired milk and blood samples from Atlantic bottlenose dolphins (*Tursiops truncatus*).^14^ Analysis of blood samples from maternal-fetal pairs reveals these trends are consistent for PFAS in the same dolphin species, as well as orcas (*Orcinus orca*).^26^ In marine mammals, it has even been noted that phospholipid concentration can be predictive of PFAS bioaccumulate in tissues, particularly the liver, kidney, heart, and brain.^27^ Since some components to make milk come from these tissues, and marine mammal milk has a higher fat content, it follows that there would be a higher mobilization of accumulated PFAS to milk. Despite this, marine ecologists, scientists, and medical professionals still lack data regarding how much neonatal PFAS exposure can be attributed to breastfeeding, what specific compounds are transferred via breastmilk, and the longitudinal trends in concentration and composition of PFAS.

In this study, we leveraged historical breastmilk samples from an individual Atlantic bottlenose over the course of a two-year, primiparous lactational period to assess trends in the maternal transfer of PFAS and draw conclusions about subsequent infant exposure. Data from this study is critical in evaluating potential threats to bottlenose dolphins, which serve not only as indicator species, but as keystone species of which the entire trophic network depends. Due to their long lifespans, extended lactation periods, and position at the top of the food chain, dolphins also serve as sentinel species for transfer of PFAS in human breastmilk. Milk, however, has historically been a difficult sample matrix for PFAS analyses. Thus, in this study we also provide an optimized PFAS extraction method for milk with varying fat contents and different volumes (1 mL or 5 mL) to allow evaluation of limited samples volumes with different fat amounts. Using a non-targeted data acquisition platform combining liquid chromatography, ion mobility spectrometry, and mass spectrometry (LC-IMS-MS), PFAS analyses were then performed in the breastmilk to inform future studies about the relationship between PFAS exposure, breastfeeding, subsequent infant health outcomes, and prevention strategies. This study therefore provides a valuable model for observing longitudinal PFAS trends and infant exposure over time—an approach that is often difficult to achieve in human studies.

## 2. MATERIALS AND METHODS

### 2.1 Sample Collection and Handling

Commercial goat’s milk purchased from Whole Foods in Chapel Hill, NC (Meyenberg; Turlock, California) was used for method development and matrix-matched calibration curves. Three quarts were combined to create a singular homogenous stock which was aliquoted into 50 mL Falcon tubes (Thermo Scientific; Waltham, MA) and stored at -20°C until use.

The Atlantic bottlenose dolphin (*Tursiops truncatus*) milk samples were obtained from a single individual, Slooper, at the Naval Command Control and Ocean Surveillance Center (San Diego, CA) (now the US Navy Marine Mammal Program), using a modified breast pump.^28, 29^ During this time, the dolphin was fed fresh frozen herring, Pacific mackerel, capelin, squid, and Columbia River smelt, which were not tested for POPs. Slooper was born at the facility in 1979 and gave birth to her first calf in 1991. Samples studied here span a gestational age for this calf from 103-706 days (1991-1993).^28, 29^ Exact sampling dates are located in **Table S1**. Samples were donated to the Smithsonian’s National Zoo and Conservation Biology Institute’s (NZCBI) Mammal Milk Repository and were stored at -20°C until analysis. All sample timepoints were deidentified prior to extraction and data analysis to eliminate potential bias.

### 2.2 Standards and Reagents

Native and stable isotope labelled (SIL) standards were purchased from Wellington Laboratories (PFAC30PAR and MPFAC-HIF-ES; Guelph, ON, Canada). All chemical reagents were purchased from Thermo Scientific (Waltham, MA), unless otherwise specified and solvents used for sample preparation and chromatography were Optima LC-MS Grade. For specific information regarding all standards, reagents, and consumables used in this work including vendor and catalog numbers, see **Table S2**.

### 2.3 Modified FDA C10.020 Extraction Method

To perform the PFAS extractions, the breastmilk samples were thawed in a water bath at room temperature and then sonicated to homogenize. PFAS was separated from a 5 mL aliquot of breastmilk using a liquid-liquid extraction based on the FDA C010.02 QuEChERS extraction and using salt pouches containing 6.0 g MgSO_4_ and 1.5 NaCl (Agilent Technologies; Santa Clara, CA). Next, two commercially available supernatant clean-up methods were tested. The first consisted of filtration through Agilent’s Captiva EMR-Lipid cartridge (3cc) after diluting to 80% with water (Santa Clara, CA). The second utilized dispersive solid-phase extraction (dSPE) containing 300 mg primary-secondary amine (PSA), 150 mg graphitized carbon black (GCB), 900 mg MgSO_4_ (Supel^™^ QuE PSA/ENVI-Carb, Millipore Sigma; Burlington, MA) with subsequent syringe filtration through 25 mm, 0.2 μm nylon filters (Whatman; Maidstone, United Kingdom). From clean-up, both extracts were concentrated to dryness using a SpeedVac, reconstituted in methanol with 4 mM ammonium acetate, and transferred to a glass LC vial with a polypropylene insert (Agilent Technologies; Santa Clara, CA). A more detailed description of these methods is shown in **Figure S1** and **S2**.

Clean-up methods were compared by calculating process efficiency for each.^30^ For the process efficiency measurements, SIL was added to a sample of goat milk prior to extraction in quadruplicate for each method. A standard blank (SB) was also prepared with neat reconstitution solvent spiked with an equal amount of SIL calculated by assuming no loss through the extraction. Process efficiency was assessed by dividing the peak areas for the 18 PFAS SIL by peak areas in the SB. Ultimately, dSPE was chosen for further extraction optimization due to its lack of extraction bias towards specific classes of PFAS which would facilitate quantitation across a greater variety of species and non-targeted analysis (NTA). Further elaboration is included in the Results section.

### 2.4 “Mini QuEChERS” Extraction

To preserve precious historical samples, a smaller version of the QuEChERS and dSPE extraction method (from 5 mL to 1 mL) was designed in an attempt to decrease all reagent volumes/masses by 80%. In brief, 1 mL of milk was added to a 15 mL tube containing 800 mg MgSO_4_ and 200 mg NaCl (UCT; Hayward, CA). Clean-up occurred as previously described, with the substitution of a 2 mL vial containing 25 mg PSA, 7.5 g GCB and 150 mg MgSO_4_ (Thermo Scientific; Waltham, MA) and syringe filtration through a 13 mm, 0.2 μm nylon filters (Whatman; Maidstone, United Kingdom). A schematic of this extraction method, hereafter referred to as “Mini QuEChERS,” is shown in **Figure S3**. This method was validated by calculating process efficiency as before.^30^

### 2.5 Dolphin Milk Analysis

Dolphin milk samples were extracted using the Mini QuEChERS method over three days, with one replicate aliquot from each sampling date per preparation batch. Additionally, a dolphin milk pool was created from all samples. One replicate of this dolphin milk pool as well as one replicate of NIST SRM 1954 (Fortified Human Breastmilk) were also extracted with each preparation batch to assess batch-batch variability. A pooled dolphin milk sample was also prepared with the calibration curve using goat’s milk on a separate day to check consistency. Both the NIST SRM and pooled samples were used as quality control (QC) samples. A method blank (MB), consisting of goat’s milk was also extracted on each sample preparation day and with the calibration curve to account for potential contamination and to later assess method detection limits (**Figure S6** and **S7**).

### 2.6 Instrumental Analysis

All extracts were analyzed with a platform that can perform targeted, suspect screening, and non-targeted analysis by interfacing liquid chromatography with drift tube ion mobility spectrometry and mass spectrometry (LC-DTIMS-MS).^31^ Exact injection order schemes are shown in **Figure S4**. Separations by retention time (RT) were conducted using an Agilent 1290 Infinity UPLC system (Agilent Technologies; Santa Clara, CA) with 4 μL injections. Reversed phase LC (RPLC) was used to determine PFAS RT as consistent with numerous prior publications.^31-35^ Briefly, each sample was injected onto an Agilent ZORBAX Plus C18 guard column (2.1 x 5 mm, 1.8 μm; Santa Clara, CA) followed by an Agilent ZORBAX Eclipse Plus C18 column (2.1 x 50 mm, 1.8 μm; Santa Clara, CA) with the column compartment held at 30°C and a flow rate of 0.4 mL per minute. Mobile Phase A consisted of 100% water, while Mobile Phase B was comprised of 95% methanol and 5% water, and both were buffered with 5 mM ammonium acetate. Exact gradients and additional information for the RPLC method can be found in **Table S3**.

IMS-MS measurements were collected using an Agilent 6560-QTOF with previously optimized methods.^31, 34^ Briefly, following ionization by the Agilent’s Jet Stream electrospray (ESI) source (Santa Clara, CA) in negative mode, ions were introduced to the drift tube with a trap fill time of 3900 μs and release time of 100 μs. The drift tube buffer gas was nitrogen (Ultra High Purity (99.999%), Airgas; Radnor, PA). Further ESI and IMS parameters are summarized in **Table S4**. MS data was collected in MS1-only mode using a quadrupole time-of-flight (QTOF) MS operating in high sensitivity mode for the *m/z* range of 50-1700.

Once each day of data analysis, ESI-L tuning mix (Agilent; Santa Clara, CA) was flow-injected with identical instrumental conditions to samples to ensure accurate calibration of the instrument for both the IMS and MS dimensions. These datafiles were also used to perform single-field calibration of observed drift times in Agilent MassHunter Workstation Software IM-MS Browser Version B.10.00 to obtain accurate and reproducible collisional cross section (CCS) values.^36^ Data files have been uploaded to MassIVE, a publicly available mass spectrometry data repository, under accession number MSV000097750. All .d files containing the RT, CCS, and MS data were evaluated using Skyline software, an open-source processing tool, which enabled suspect screening against an in-house multidimensional library.^37^ All identifications had RTs within one minute of in-house library standards, mass error ≤ 10 ppm, and CCS values within the resolving power window of 30.

### 2.6 Quantitative Analysis

A matrix-matched calibration curve was prepared in commercial goat’s milk. The 10-point calibration curve (0.01 to 50 ng/mL) was extracted alongside the dolphin milk samples and QCs. To quantify, matching SIL standards were used to calculate light-to-heavy peak areas, and in cases where a direct SIL match was not available in the standard mix used, a surrogate from a similar PFAS class was used (see **Table S5** for exact analytes used for normalization). The method detection limit (MDL) was defined based on guidance from EPA 821-R-16-006, or the concentration of analytes in the method blanks (**Figure S8**).^38^ The limit of quantitation (LOQ) was defined as the lowest point on the calibration curve where the relative standard deviation (RSD) of injection replicates was less than 20%.^39^ All calibration curves were fit such that R^2^ ≥ 0.98 with 1/(x*x) regression weighting. When branched isomers were identified, they were integrated with the linear to calculate a total concentration for all isomers. Later, these were integrated separately, and peak areas were used to determine the ratio of branched to linear isomer.

### 2.7 Suspect Screening and Non-targeted Data Analysis

Using Skyline software and a publicly available in-house library containing CCS values, RT, and *m/z* built using certified standards, suspect screening analyses were conducted for 156 PFAS.^37^ For the suspect screening and non-targeted data analysis where there are no calibration standards available to compare concentrations relative to the method blank, and the MDL was based on the raw peak area of the method blanks, rather than concentration (**Figure S8**). To further validate isomeric structures for some PFAS species, travelling wave ion mobility (TWIMS) with MOBILIon SLIM was employed (**Table S6**).

## 3. RESULTS AND DISCUSSION

### 3.1 Extraction Method Optimization

PFAS analysis for milk and other dairy products has historically proven difficult. The United States Food and Drug Administration’s (FDA) has previously published generalized methods for the determination of PFAS from food products, C-010.03, which employs a QuEChERS extraction followed by optional dSPE cleanup. Despite the publication of this original version of the method in 2019, most breastmilk studies in the last 5 years employ ion-pairing extraction, weak anion exchange, protein precipitation, or traditional liquid-liquid extraction methods to isolate PFAS from human breastmilk.^40, 41^ In some applications, these are combined with subsequent sample clean up with solid phase extraction (SPE), which is common for analyses of drinking water.^42-46^ While these methods have proven useful for other matrices, breastmilk is more compositionally similar to food, suggesting that the FDA’s QuEChERS method may be more appropriate. Only one group (Olowojo) has adopted the method proposed by the FDA for human breastmilk analysis.^47^ Furthermore, this method has only been adopted for studies of human breastmilk, which are often not limited by sample volumes. Additionally, most studies have only performed targeted quantitative PFAS analyses and not looked for new and unknown PFAS. Since the goal of NTA is to perform a largely unbiased extraction to study as many PFAS as possible, regardless of head group functionalization, methods capable of extracting a large range of compounds are needed.

Dolphin milk also possesses significantly higher fat content (13-18% fat)^48^ in comparison to human or dairy milk (3-5% fat), for which these methods were originally developed.^49,50^ Thus, we sought to evaluate the efficiency of variations on the QuEChERS method to extract PFAS from four classes in milk samples with varying fat contents and lower sample volumes than previously described in the literature.

Since the QuEChERS method requires a supernatant clean-up step following biphasic extraction, we compared two different methods including the carbon-based dSPE, and Agilent’s Captiva EMR Lipid cartridge which is designed to specifically remove lipids which may otherwise hinder PFAS analyses (**Figure 1A**). To compare these two cleanup strategies, we calculated the average process efficiency (PE) for 18 SIL PFAS across four classes: perfluoroalkyl carboxylic acids (PFCAs), perfluoroalkyl sulfonic acids (PFSAs), perfluoroalkyl sulfonamides (PFASAs) and fluorotelomer sulfonic acids (FTSs) spiked in milk. PE can generally be thought of as a combination of analyte recovery throughout the extraction process which also accounts for matrix effects including ion suppression (**Figure 1B**). A PE of 100% is ideal, showing optimal extraction and ionization of each analyte in the different matrices. In our study, the dSPE clean- up method outperformed EMR as evidenced by consistent PEs near 100% for all analyzed PFAS classes (**Figure 1B**). This indicates that dSPE does not bias extraction towards specific PFAS classes and matrix effects do not impede efficient ionization of PFAS species. It is likely that the greater losses for the PFCAs and the PFASAs with the EMR method are due to their headgroup similarities to those of lipids, resulting in unwanted preferential binding to the sorbent’s stationary phase. Of note, the FTSs and PFASAs have a PE greater than 100% with EMR since the headgroups are less attracted to the stationary phase and therefore are transferred more readily to the final extract. Based on these results, and data that dSPE had more consistent PEs around 100% for all classes evaluated, QuEChERS with dSPE cleanup was selected for further optimization.

**Figure 1.**
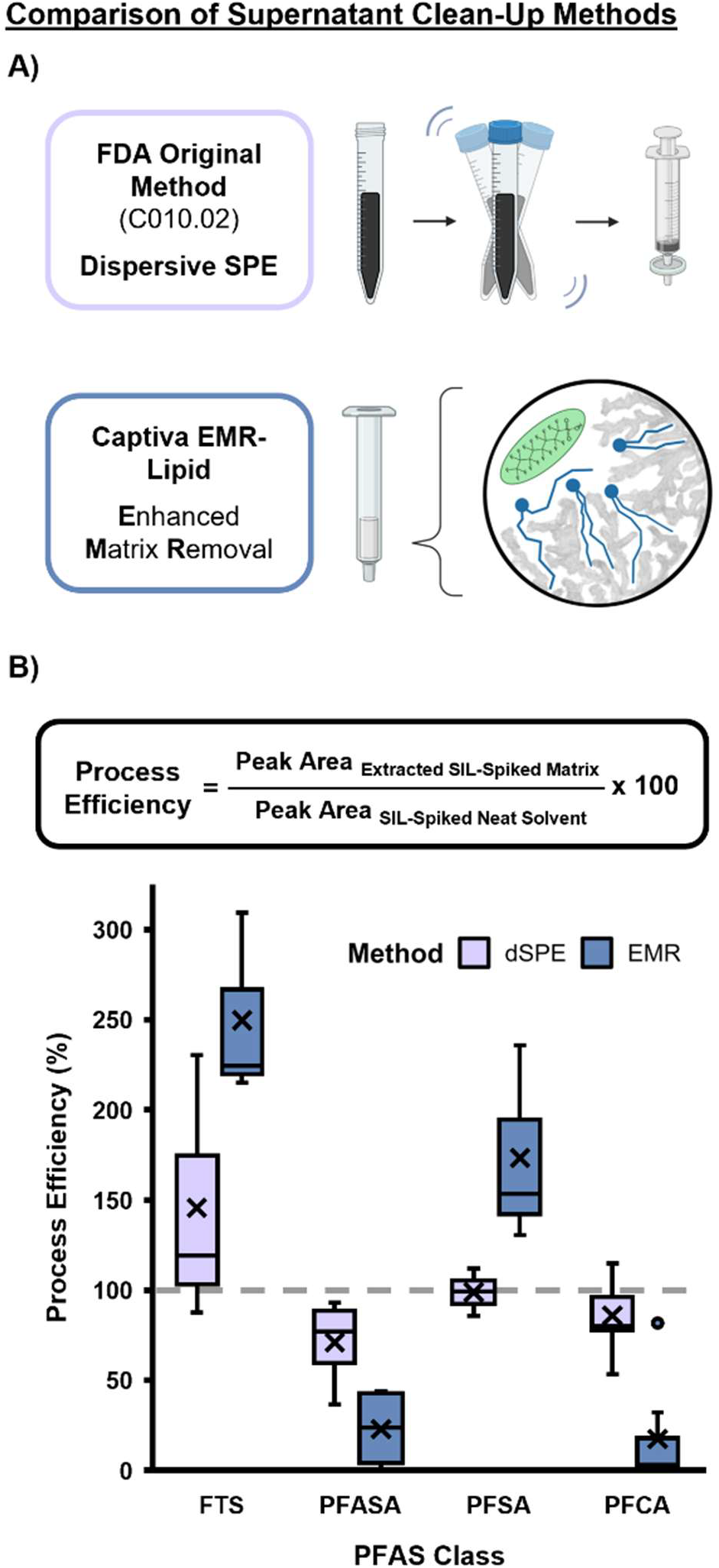
A Comparison of PFAS Extraction Clean-Up Methods for Milk. **A)** This study evaluated the FDA’s suggested method involving addition of supernatant to a dSPE tube containing graphitized carbon, primary-secondary amine and MgSO_4_ and syringe filtration, as well as Captiva Enhanced Matrix Removal (EMR) Lipid cartridge. **B)** Methods were evaluated by calculating PEs based on spiked SIL peak areas. The dSPE method showed PEs centered around 100%, while EMR shows PE varying based on class. Boxes represent interquartile range (IQR), with whiskers extending to 1.5 IQR and individual points representing outliers. X’s represent mean.

In general, breastmilk is a precious resource, and we realize 5 mL may be difficult to obtain in some studies. Thus, we sought to reduce the initial starting volume of milk from 5 mL to 1 mL using the QuEChERS extraction with dSPE cleanup, through the development of a “Mini QuEChERS” approach. We feel this miniaturization is also important in preserving historical samples which may be useful for later POP analysis. To achieve this, we reduced the reagent volumes, QuEChERS and dSPE salt masses and consumable sizes by approximately 80% but kept the normal QuEChERS workflow (**Figure S3**). Interestingly, the calculated PEs for the Mini QuEChERS method were comparable, or in some cases, higher than that of the original method (**Figure 2A**). This may be due to slight differences between reagent manufacturing processes, or a result of lower volumes being transferred, which reduces errors in the extraction. N-methylperfluorooctane sulfonamidoacetic acid (NMeFOSAA) has a PE much higher than 100% for the Mini QuEChERS method even though the original falls below 100%, which may be attributable to analyte specific solubility at lower sample volume or manufacturer variation in QuEChERS materials. In addition to comparable PE performance, the reduction in sample volumes still resulted in low LOQs and MDLs (**Figure 2B**) without unreasonable variation between different species or classes. All LOQs and MDLs were below 1 ng/mL, with most LOQs below 0.1 ng/mL. For all the PFASAs, there was zero signal detected in the method blanks, thus the MDL was defined as any appreciable signal following demultiplexing.

**Figure 2.**
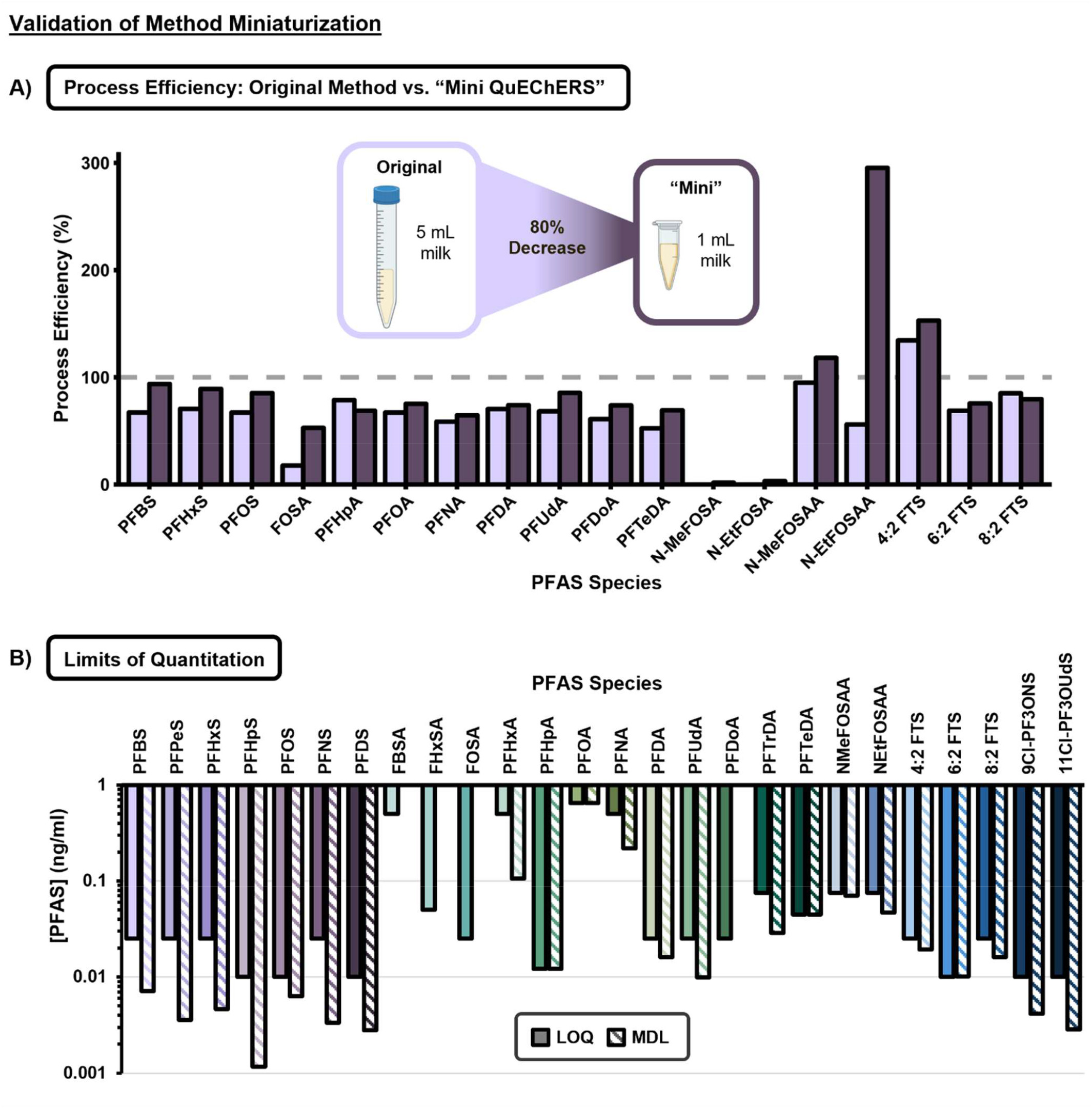
Validations of “Mini QuEChERS” PFAS Extraction from Milk. **A)** PE was used to compare the original QuEChERS method with dSPE using 5 mL starting volume to a miniaturized method named “Mini QuEChERS” with 80% decrease in method materials and starting volume (1 mL). The miniaturized method was found to have comparable or higher PE for all analytes when prepared in quadruplicate. **B)** Method detection limits (MDL) and limits of quantitation (LOQ) were calculated based on quadruplicate injection replicates of a matrix-matched calibration curve in goat’s milk. In cases where no MDL is shown, the MDL is defined as showing any appreciable signal after demultiplexing.

### 3.2 Longitudinal Trends in Lactational PFAS Transfer

Previous studies have demonstrated that the lactational burden of POPs decreases following birth. To determine if PFAS follow a similar trend, we analyzed the milk from a single dolphin over the course of a two-year lactation for her first calf. This analysis included 21 sampling dates over the 2 years which were on average 30 days apart, giving a rough estimate of month-month variation of PFAS in the milk (**Table S1**). Based on the samples provided by the NZCBI, the earliest sample analyzed corresponds with an infant age of 103 days, or approximately three months old. Additionally, for each sampling date, three separate 1-mL aliquots were analyzed to account for variation in the milk sampling and biological variability over a feeding session.

In total, 26 PFAS species were targeted for quantification in this study (**Table 1**) based on constructed calibration curves. Of these, 13 were detected above the LOQ in at least 2/3 replicates from at least one sampling date, the threshold for identification established prior to analysis (**Table 1**). In addition to high concentrations of PFSAs and PFCAs (>1 ng/mL), PFAS from the FTSs, and PFASAs were also detected at quantifiable concentrations, demonstrating that milk can expose infants to a variety of PFAS regardless of chemical moiety. Specifically, the legacy PFAS, perfluorooctane sulfonic acid (PFOS) was found to be the greatest contributor to total PFAS in milk across all sampling dates, with the highest average concentration of 7.18 ng/mL in a sample at the earliest date (infant age equal to 103 days) (**Figure 3A**). The longitudinal analyses illustrate that the concentration of PFOS generally decreases over time. These findings are consistent with previous studies evaluating trends for common organochlorine compounds such as polychlorinated biphenyls (PCB) and dichlorodiphenyldichloroethylene (*p*,*p’*-DDE), for the same dolphin milk samples as reported by Ridgway and Reddy.^28^

**Table 1.**
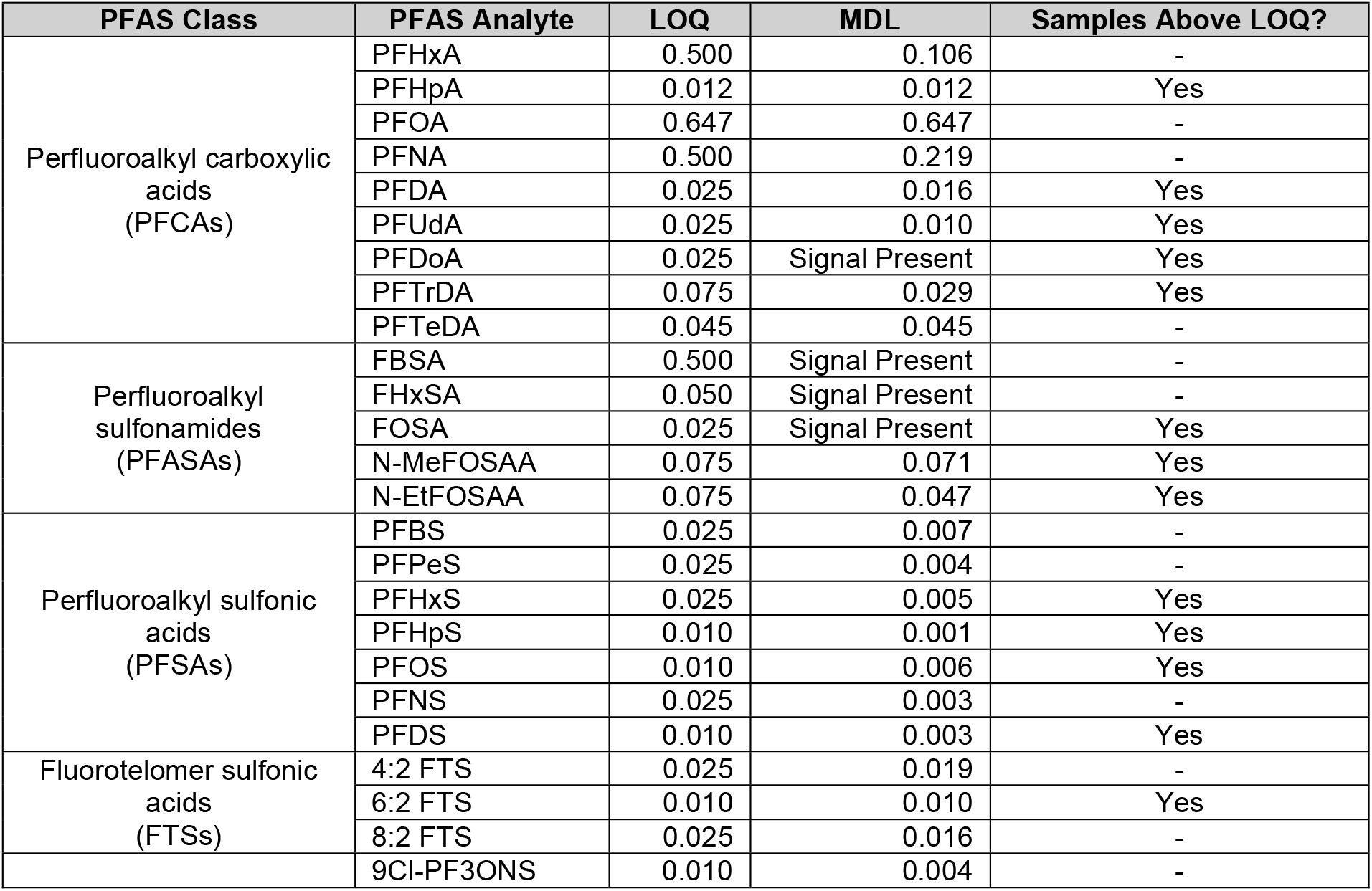
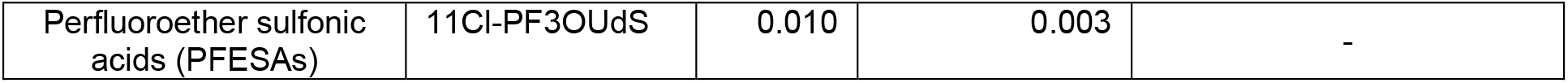
PFAS Targeted for Quantitation, including MDLs and LOQs. 26 PFAS species were targeted for quantitation, with 13 being above the LOQ in at least 2/3 replicates for at least date. Analytes which were not detected above the LOQ were later analyzed for detection frequency above the MDL based on concentration (**Figure S8**).

**Figure 3.**
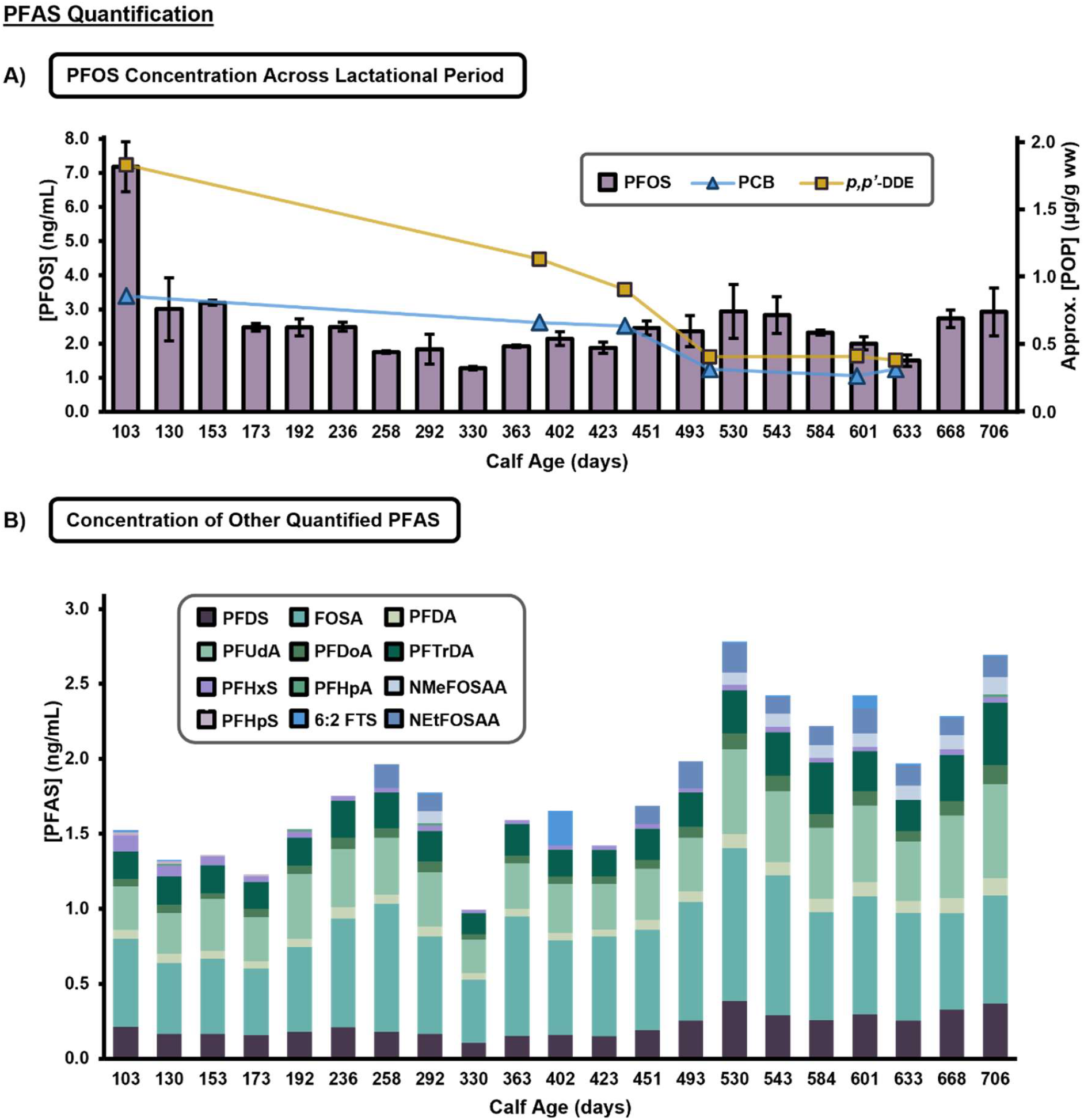
Quantitative Longitudinal Trends in PFAS found in Dolphin Milk. **A)** Average PFOS concentrations (ng/mL) at each sampling date are shown by the purple bars and left y-axis, with the highest average exposure at the earliest date (103 days). Data from previous studies evaluating trends for common organochlorine compounds PCB and *p*,*p’*-DDE, as reported for the same dolphin milk samples by Ridgway and Reddy, are overlayed and display similar longitudinal trends as shown by the blue and yellow line and right y-axis.^28^ Error bars represent the standard deviation of each sample. **B)** Average concentrations (ng/mL) for 12 additionally quantified PFAS representing four PFAS classes at each sampling date.

After PFOS, long-chained PFAS (≥C8 compounds), such as perfluoroundecanoic acid (PFUdA, C11), perfluorododecanoic acid (PFDoA, C12), and perfluorotridecanoic acid (PFTrDA, C13), had the next highest concentrations found in dolphin milk throughout the lactation (**Figure 3B**). Previous analyses of marine mammal tissue and blood also reported higher proportions of long-chained PFAS, and this finding further exemplifies that long-chained compounds are a major type of exposure from breastmilk.^14, 27^ Additionally, studies of whales have illustrated that tissues with higher relative phospholipid concentrations, like the liver, are associated with higher concentrations of PFCAs accumulating in those tissues, and this trend was exacerbated for longer-chained PFAS.^27, 51^ Based on this evidence, we hypothesize that PFAS accumulated in these fattier tissues and the liver is mobilized to the mammary glands during milk production. This would not only account for the higher proportion of long-chain PFAS found in milk, but also the longitudinal trends observed for the total PFAS concentration as the body relies on stored energy sources when demand for calories and nutrients is high at the beginning of lactation. Therefore, bioaccumulated PFAS likely contributes to the overall concentration of PFAS in dolphin milk, reducing maternal body burden, but increasing exposure for first-born calves early in the lactational period.

It is important to note that Atlantic bottlenose dolphins, and cetaceans more broadly, are known to predominantly rely on increased food consumption to offset the energetic cost of lactation.^52-54^ Numerous studies have indicated that fish accumulate PFAS and are a major source of dietary PFAS exposure.^55-58^ Therefore, it is likely that a large portion of the PFAS observed in the milk comes from diet, especially later in lactation. This study found the PFASAs, NMeFOSAA and N-ethyl perfluorooctane sulfonamido acetic acid (NEtFOSAA), increase in abundance towards the end of the sampling period while 6:2 FTS levels vary throughout the lactation period. Together, this may be indicative of a point-source exposure or changes in food sources. Designed longitudinal studies which evaluate milk, as well as maternal and neonatal blood, could identify changes in circulating PFAS to determine the differential contribution from bioaccumulated PFAS versus diet.

It is widely accepted that high concentration of PFAS can induce adverse health outcomes in exposed populations. In marine mammals, PFAS are known to accumulate preferentially in fatty tissues, like the brain and liver,^27, 59^ so they may also induce adverse health effects in exposed calves at crucial stages of growth and development.^26^ Also troubling, a neonatal dolphin’s blubber layer thickness increases rapidly following birth.^60^ If that dolphin is concurrently receiving high doses of PFAS through breastfeeding, the materials from the milk used to support blubber layer growth are likely leading to accumulation of PFAS in the neonate. To further contextualize the amount of PFAS to which the dolphin calf in this study was exposed, we utilized, consumption data from a hand-reared dolphin at three months of age (**Figure S10**).^61^ Using this data, the dolphin calf around 103 days old would intake 4,200 mL of milk per day, or 29,400 mL per week. Since we observed 7.18 ng of PFOS per milliliter of milk, this corresponds to roughly 5,300 nanograms per kilogram bodyweight per week (ng/kg bw/week). When taking all quantified PFAS species into account, this exposure increases to 6,500 ng/kg bw/week (**Figure 4**). In stark contrast, the regulatory agency, Food Standards Australia New Zealand (FSANZ), has set a health-based guidance value for PFOS at merely 140 ng/kg bw/week for humans based on toxicological laboratory animal studies.^62^ The European Food Safety Authority’s guidance is even more conservative, at only 4.4 ng/kg bw/week for PFOS in healthy adult humans.^63^ Since fish are recognized by both agencies as a primary dietary source of PFAS, the elevated exposure in dolphin calves expected, however this great extent of PFAS exposure remains troubling, especially as the guidelines do not consider additional PFAS species. Additionally, our longitudinal trend suggests that even higher PFAS exposure will likely occur before the earliest sampling date presented in this work. Since calf mortality in dolphins peaks in the first 30 days of life,^64^ high levels of exposure may worsen these outcomes or have other long-term health effects. In other mammals, daily PFAS intake is correlated with higher circulating PFAS levels in infancy.^65^ In a different study of wild bottlenose dolphin plasma, it was found that chronic exposure to PFCAs was related to alteration of immune system function and markers of liver damage.^23^ These findings suggest that elevated PFAS concentrations in milk throughout infancy may lead to other concerning outcomes, and further research is needed to fully understand the lifelong impacts of lactational exposure.

**Figure 4.**
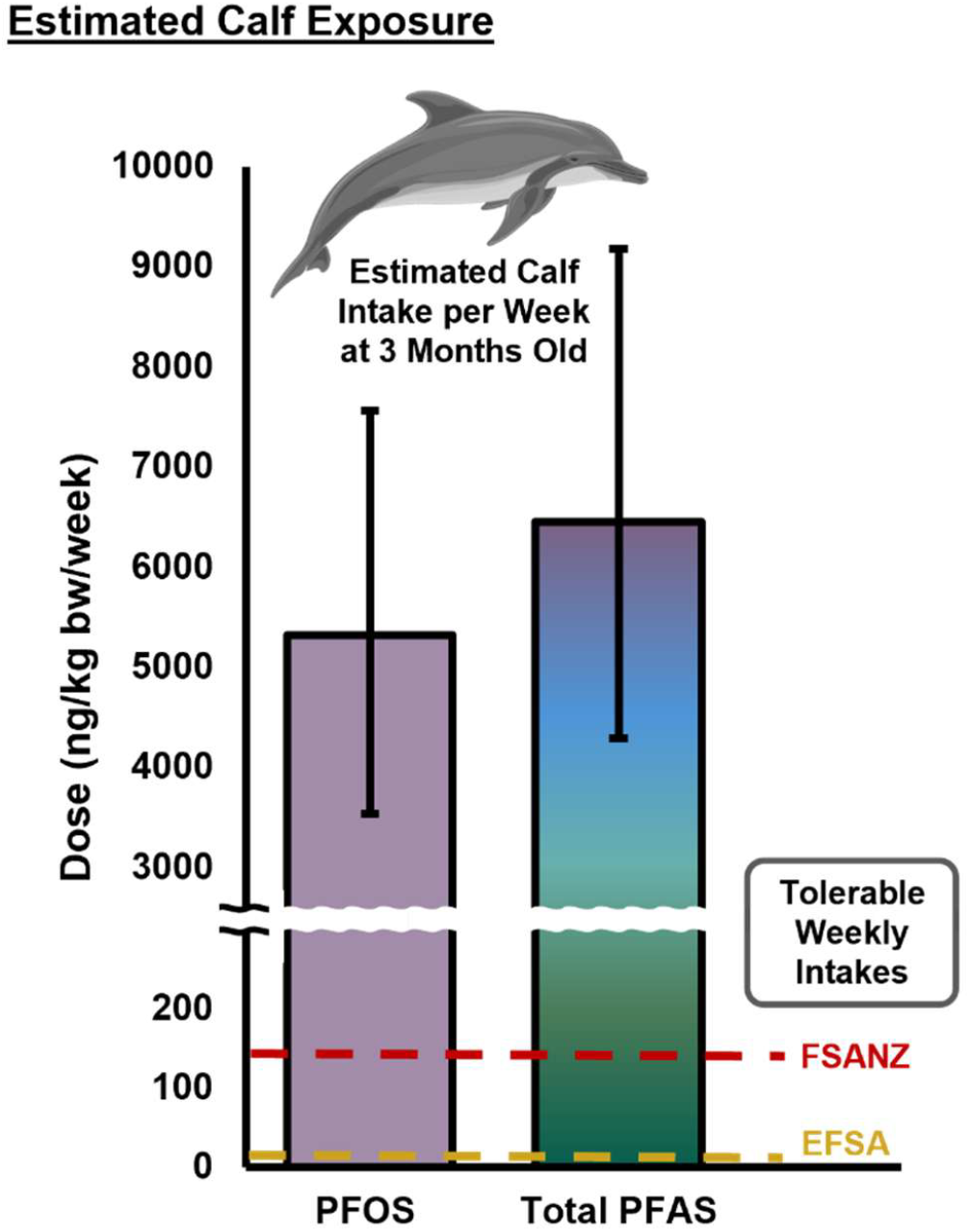
Estimated Calf Exposure to PFAS Concentrations at 3 Months of Age. Based on the estimations of daily consumption of milk and the weight of an Atlantic Bottlenose Dolphin, it is likely that the dose in ng PFAS per kilogram bodyweight per week at approximately 3 months of age (PFOS ≈ 5300 ng/kg bw/week, ∑PFAS ≈ 6500 ng/kg bw/week) far exceeds recommended tolerable weekly intakes as published by Food Safety Australia New Zealand (FSANZ, 140 ng/kg bw/week, PFOS) and the European Food Safey Authority (EFSA, 4.4 ng/kg bw/week, PFOS). Error bars represent lower and upper estimations based on published values

### 3.3 Neonatal Exposure to Emerging PFAS

Although research on neonatal PFAS exposure is expanding, many epidemiological studies examine only one compound, failing to capture the complexity of mixtures encountered in the environment. Here, we conducted suspect screening and NTA with IMS to allow for a more extensive evaluation of fluorinated features found in dolphin milk samples. Each sample is screened against a library containing RT, CCS values, and *m/z* ratios, often for multiple ion types, for 156 PFAS species. The multidimensional analysis allows for expanded scope of PFAS analysis, isomeric separation, and more confident PFAS identifications.^31, 33, 37, 66^ For some PFAS, multiple peaks were observed, many of which were unresolved. Upon further investigation using the MOBILion MOBIE 2.0 platform (EyeON version 2.3) which has an IMS resolving power of ∼200, it was confirmed that the earlier eluting peaks correspond with different mobility profiles, suggestive of branched isomers (**Figure 5A**). This phenomenon was also reported in Dodds *et al*. (2020), where IMS facilitated the separation of branched isomers which traditional LC-MS/MS methods may have difficulty characterizing.^31^ While very few studies look at the effects of linear and branched structures separately, they are known to have variable accumulation in biological matrices, where the branched forms may be less potent due to lower protein-binding affinity.^67-69^ However these studies are limited to perfluorooctanoic acid (PFOA) and PFOS, and related isoforms. In this study, branched isomers of PFSAs (C6 to C10), PFCAs (C11 to C13) and one PFASA were detected in dolphin milk for multiple sampling dates (**Figure 5B**). The median ratio of branched:linear isomers ranged from 1 to 15%, indicating that branched structures may make up a sizable proportion of the quantified PFAS in this work. These branched structures may also have varying biological impacts based on their ability to accumulate or act in metabolic pathways, highlighting the utility of multidimensional separations for PFAS NTA.

**Figure 5.**
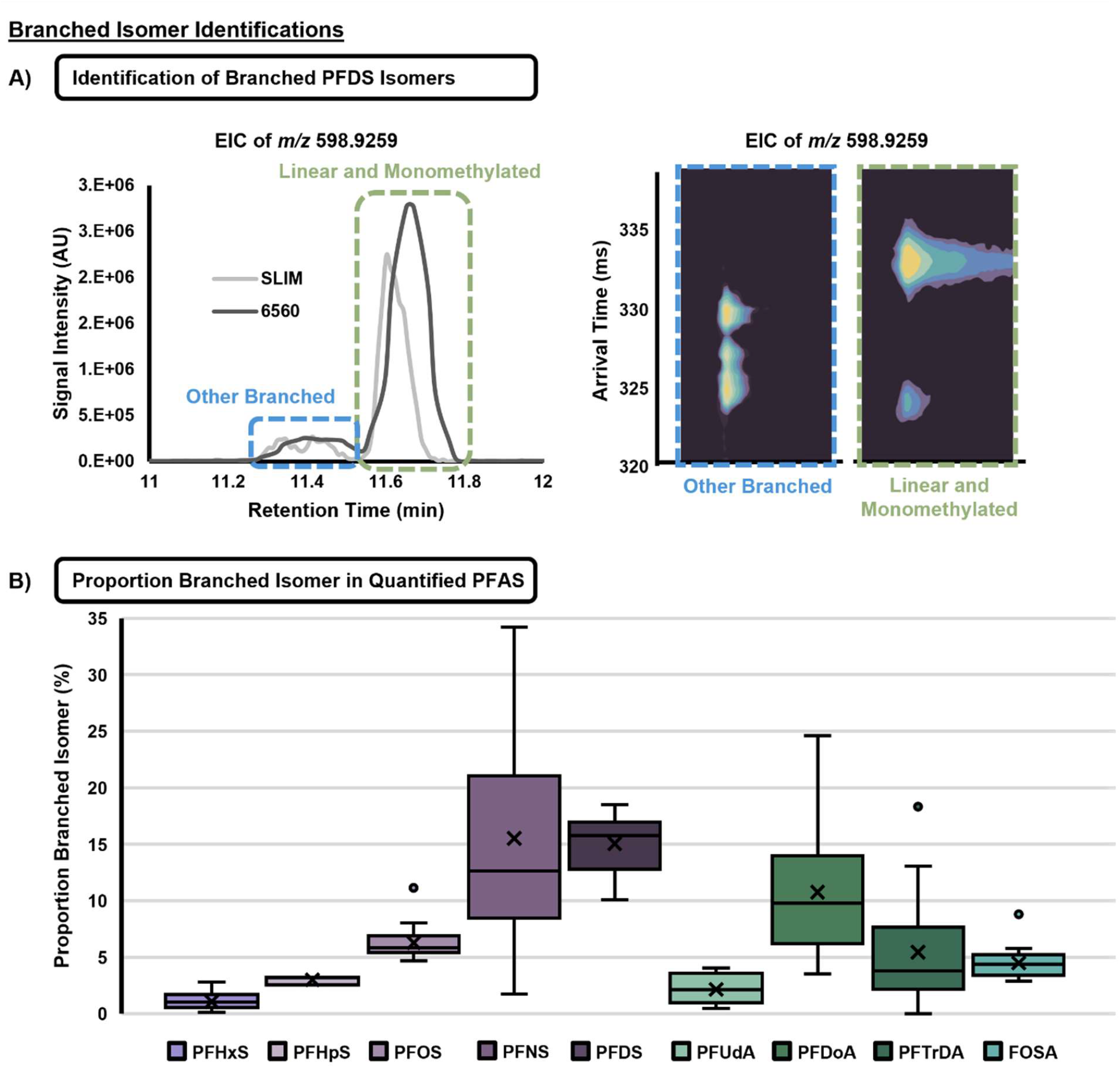
Branched Isomer Identifications. **A)** Extracted ion chromatograms for PFDS collected using the MOBILion MOBIE 2.0, to show that earlier eluting peaks correspond with different mobility profiles, suggestive of branched isomers based on the different arrival times. **B)** Analysis of the ratio of branched isomer peak area to linear peak area (that which corresponds with library CCS values), several species of quantified PFAS had branched isomers, ranging from median 1.0 to 12.6% branched. Boxes represent interquartile range (IQR), with whiskers extending to 1.5 IQR and individual points representing outliers. The X’s on the bars represent mean values.

Although no distinct temporal trends were observed in the proportion of branched isomers (**Figure S11**), results indicate that carbon chain length increases for PFSAs with proportion of branched isomer in the milk (**Figure 5B**). Another study reported similar findings, noting that an increase in carbon chain length from PFHxS to PFOS was associated with greater postpartum maternal transfer of the branched isomer.^70^ However, this trend was not observed for the PFCAs in either case. Thus, it follows that both chain length and head group functionality may factor into the differential transfer of various PFAS isomers via lactation.

In addition to annotating PFAS isoforms, the multidimensional NTA approach used here allowed for suspect screening for additional PFAS based on an in-house LC-IMS-MS library.^37^ Such analyses facilitated the annotation of 13 additional PFAS which would otherwise be missed in traditional targeted LC-MS/MS screening which are noted in **Figure 6**. All identified compounds were assigned an identification confidence level based on guidance by Boatman *et al* and these are noted in **Table S7**.^71^ Furthermore, two of these compounds, perfluoroundecanesulfonic acid (PFUnS, C11) and perfluorotetradecane sulfonic acid (PFTrS, C13), were not included in the suspect library, but upon identification of the C10 and C12 PFSAs, RT vs. *m/z* and CCS vs. *m/z* trendlines were used to identify and validate these structures, a major benefit of including IMS analyses (**Figure S12**). Interestingly, three of the suspect screening compounds, PFUnS, perfluorododecane sulfonic acid (PFDoS) and SAmPAP, were found to be present throughout the lactation course. SAmPAP, predominantly used in paper coatings until the early 2000s, is frequently detected in marine sediments, and is broadly recognized to be a threat to aquatic biota.^72, 73^ It is also a degradation precursor to PFOS, and transformation studies suggests that the presence of this compound could contribute to further accumulation of PFOS in aquatic organisms.^73^

**Figure 6.**
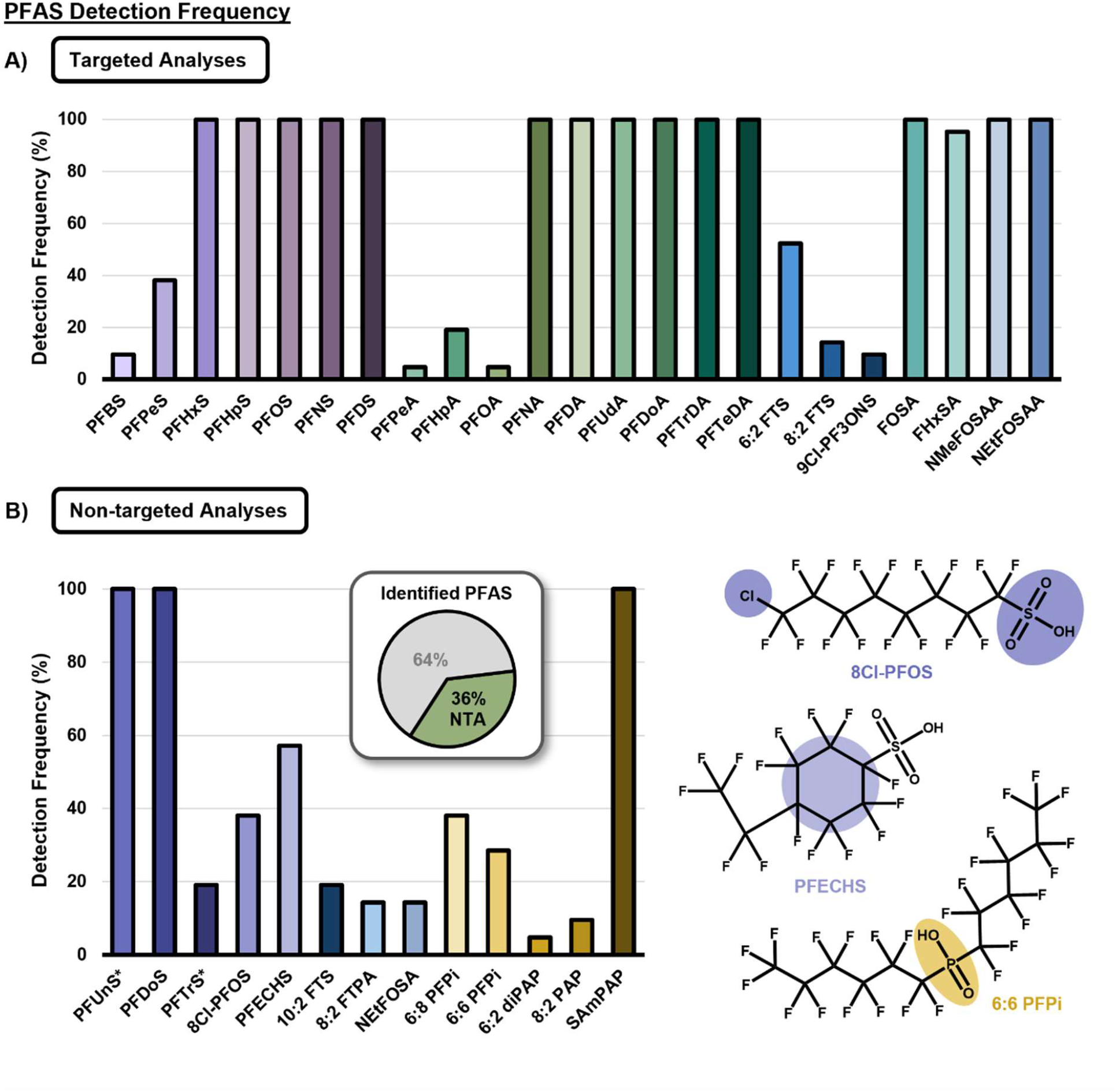
PFAS Detection Frequency for Targeted and Non-Targeted Analyses. **A)** Detection frequency for targeted analytes was determined based on MDL (concentration) (**Figure S8**) and presence in at least 2/3 extraction replicates for each sampling date. **B)** Suspect screening analytes account for 36% of all PFAS detected. Example structures for 8Cl-PFOS, PFECHS, and 6:6 PFPi are shown as representative emerging/replacement PFAS detected. (*) indicates that PFAS were added by observation and validated using RT and CCS vs. *m/z* trendlines.

Suspect screening also highlighted the presence of multiple representatives from the perfluoroalkyl phosphinic acid (PFPiA) class. PFPiAs are used as surfactants, antimicrobial products, and stain repellants for carpets and upholstery.^74^ They have a high potential for bioaccumulation in fish but lack sufficient data regarding acute hazards to aquatic life.^74^ We also identified several replacement PFAS, including PFECHS and 8Cl-PFOS, which are used to inhibit erosion in hydraulic systems and aqueous film-forming foams (**Figure 6B**).^75, 76^ Although the dolphin’s exposure to these compounds may be due to its proximity to active military operations and bases, these compounds are widespread and thus, their exact source cannot be determined.

The number and variability of PFAS identified in this study, including 23 from targeted analyses and 13 from suspect screening and NTA, highlights the need for further analysis of mammalian milks. Specifically, we need to understand if these trends are specific to dolphins, or if they hold true for multiple species, regardless of ecosystem or place on the food chain. It is likely that in dolphins, PFAS in milk comes both from diet and maternal tissue where it has bioaccumulated. The results presented here therefore serve as a warning sign that neonatal mammals may be exposed via milk to a complex mixture of PFAS at high concentrations and species at higher trophic levels, crucial to ecosystem health and sustainability, are likely most at risk.

## 4. CONCLUSIONS

Milk is an essential substance for transferring energy, nutrients, and biochemical signals from a mother to their infant, however, we have identified lactation also serves as a major source of exposure to PFAS. Using a newly optimized QuEChERS extraction method for small sample volumes, we were able to extract a variety of PFAS chemical moieties from milk with a high fat content. A total of 36 PFAS were detected at least once during the lactational period, with 17 observed during all sampling dates.

Quantitative analyses revealed that dolphin milk samples across a two-year lactational period correspond with exposure to neonatal dolphin calves surpassing the recommended dietary intake based on toxicological and epidemiological studies by >1000-fold. It is clear that marine mammals are a highly exposed population, and continued exposure could have significant ramifications throughout the trophic network. Although previous studies have linked traditional legacy PFAS, PFOS and PFOA, to adverse outcomes in dolphins and other marine mammals, there remains virtually no data on the impact of these chemicals or their replacement compounds on growth and development of neonatal marine mammals, especially with dosages of this magnitude. While dolphins serve as a sentinel species for human exposure, variations in internal energy storage, milk composition and fat percentage, as well as life history may impact the varieties and concentrations of PFAS transferred via lactation between species. However, it is likely that similar longitudinal and accumulative trends presented here are conserved across species. Nonetheless, this study exemplifies the need for future studies evaluating the rate of maternal PFAS transfer by lactation and those assessing potential adverse health outcomes in all mammals.

It is important to note that the samples in this study were collected during the 1990s, prior to phase-outs of legacy PFAS compounds. Through the implementation of IMS, we were able to obtain enhanced isomeric information for these quantified legacy PFAS, where branched structures account for up to 15% of the quantified concentration, which may behave differently in the body than more commonly targeted linear isomers. The detection of SAmPAP and 12 other interesting emerging PFAS in this study through suspect screening highlights the need for NTA for PFAS studies of complex biological matrices. Even though these pollutants persist in the environment, with over 7 million potential PFAS structures in existence, strategies to detect emerging compounds are therefore more important than ever.

## AUTHOR mINFORMATION

## Corresponding Author

Erin S. Baker, *Department of Chemistry, University of North Carolina at Chapel Hill, Chapel Hill, NC 27514*

## Funding Sources

This work was funded by a grant from the National Institute of Environmental Health Sciences (P42 ES027704) and a cooperative agreement with the Environmental Protection Agency (STAR RD 84003201). The views expressed in this manuscript do not reflect those of the funding agencies.

## Notes

The authors declare no competing financial interest.

## ACKNOWLEDGEMENTS

We would like to thank Dr. Mark Strynar for his continued guidance in PFAS analyses and the generosity of the Smithsonian’s National Zoo and Conservation Biology Institute for their support of this work. Special thanks to Emily C. Vincent for her assistance in creating figures and handling strange milk. The views expressed in this manuscript do not reflect those of the funding agencies.

## Notes

### Competing Interest Statement

The authors have declared no competing interest.

