## Supplemental Information for "Maternal PFAS Transfer through Lactation: Dolphin Milk Reveals Routes of Early-Life Exposure"

### Table of Contents

Cover page and table of contents ........................................................................................ **S1-2**

**Figure S1.** QuEChERS Liquid-Liquid Extraction Scheme......................................................... **S3**

**Figure S2.** Supernatant Clean-up Method Schemes.................................................................... **S4**

**Figure S3.** Mini QuEChERS Full Extraction Scheme ................................................................ **S5**

**Figure S4.** Dolphin Milk Instrumental Analysis Workflow ....................................................... **S6**

**Figure S5.** Comparison of Quantitation Normalization Methods ............................................... **S7**

**Figure S6.** Analysis of Preparation Batch Variability with Pooled Dolphin Sample.................. **S8**

**Figure S7.** Determination of Accuracy to NIST SRM 1954 ....................................................... **S9**

**Figure S8.** Definitions of MDL and LOQ for Quantitative and Non-targeted Analyses .......... **S10**

**Figure S9.** Individual Quantified PFAS by Sampling Date ................................................ **S11-14**

**Figure S10.** Example of Dose Estimate Calculations (PFOS) .................................................. **S15**

**Figure S11.** Branched Percentage of Quantified PFAS by Sampling Date ......................... **S16-17**

**Figure S12.** NTA Identification of Long-Chain PFSAs by Trendline....................................... **S18**

**Figure S1.** A representative scheme for the original QuEChERS liquid-liquid extraction.

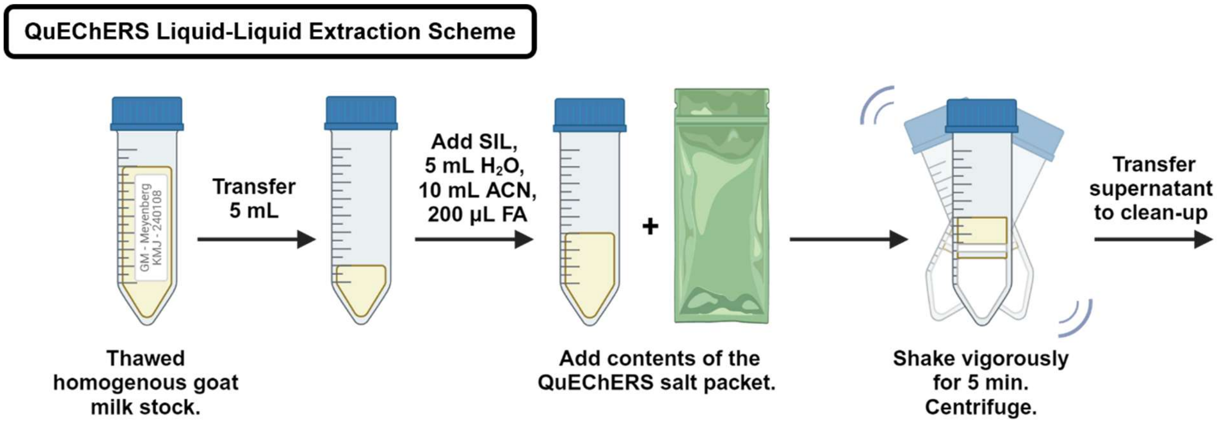

**Figure S2.** Schemes for supernatant clean-up methods: Dispersive SPE Method derived from FDA Method C010.03 and Captiva EMR-Lipid.

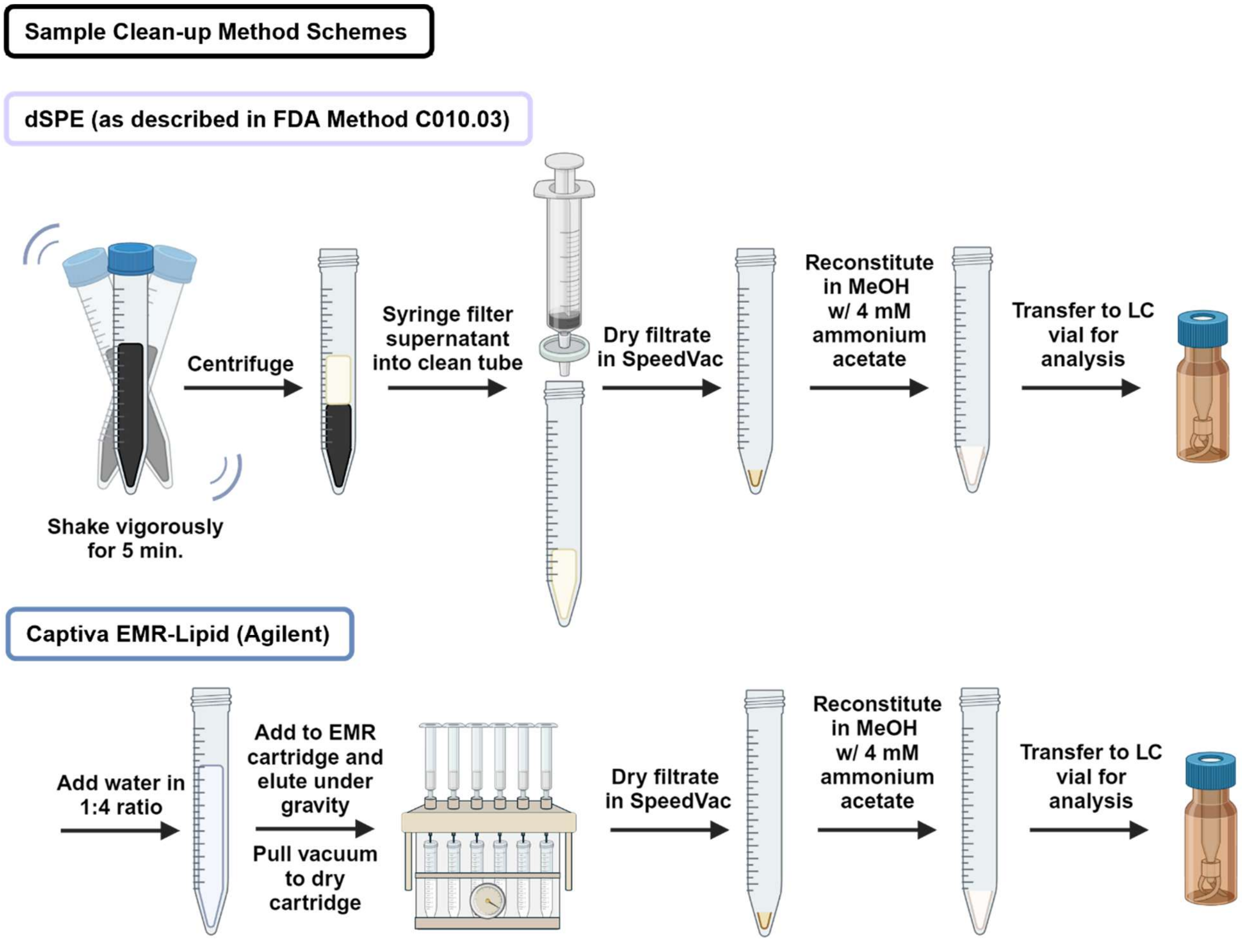

**Figure S3.** Representative scheme for the full optimized “Mini QuEChERS” PFAS extraction method used for sample analysis.

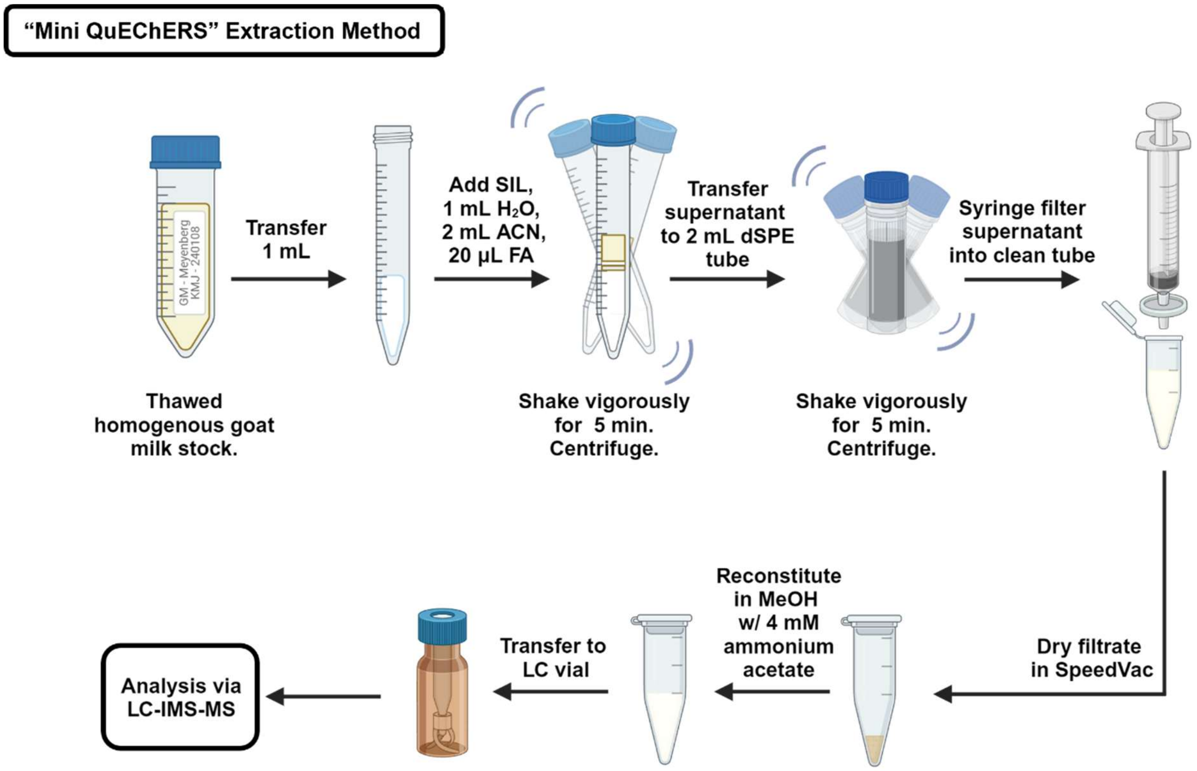

**Figure S4.** Example well-play layout for dolphin sample analysis, including dolphin milk extracts, quality controls (NIST 1954 Fortified Human Breastmilk (n=3) and pooled dolphin sample (n=4)), calibration curve and necessary blanks. Calibration curve standards were injected from low to high, once at the beginning of the worklist and again after every 1/3 of the unknown dolphin samples. Double blanks (neat reconstitution solvent) and standard blanks (neat reconstitution solvent spiked with known concentrations of native and SIL standards) were run repeatedly throughout the worklist to assess column carryover and instrument performance.

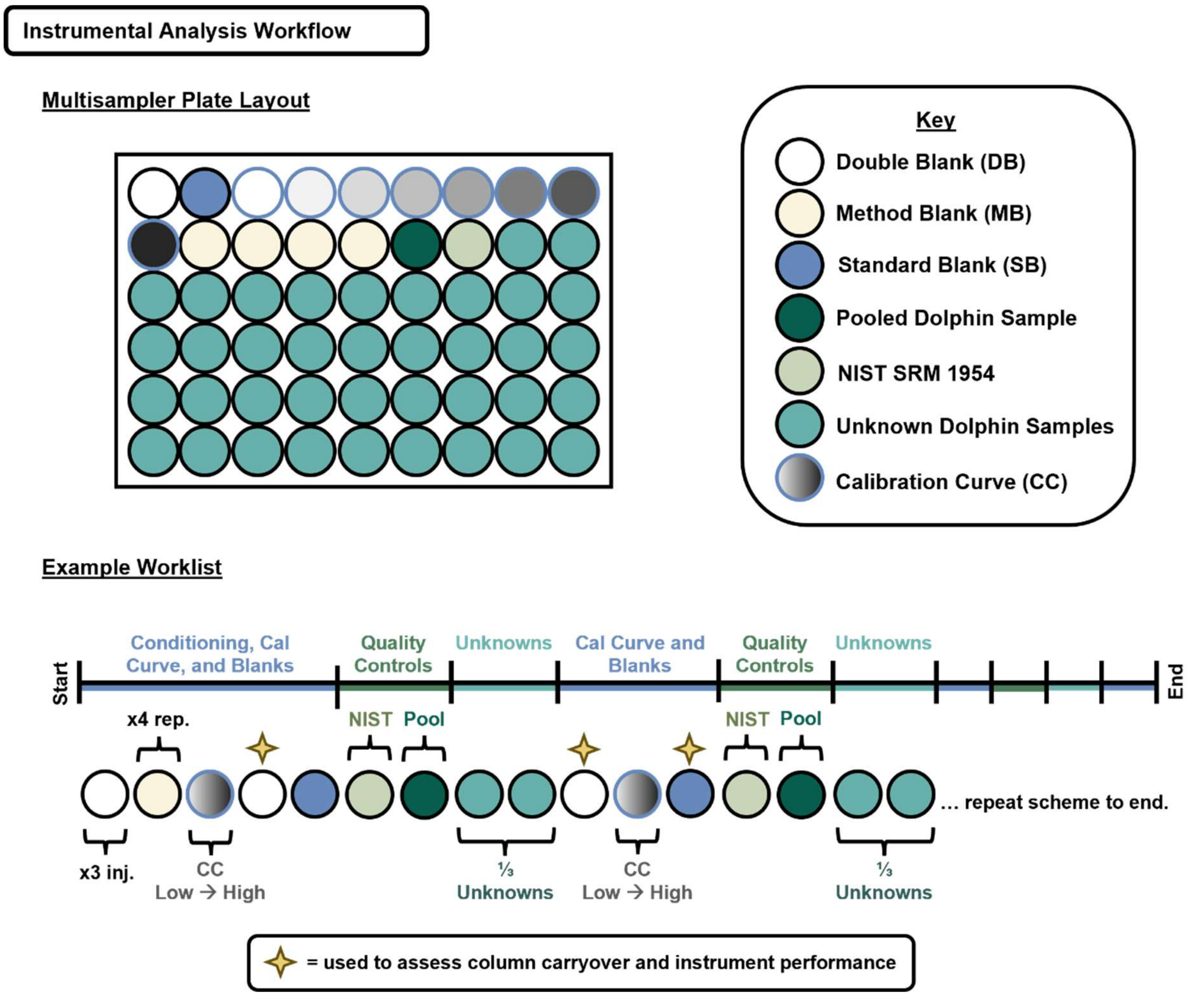

**Figure S5.** Comparison of quantitation normalization methods, either to ng/mL or ng/g ww milk for PFOS as a representative analyte. There was no difference in normalization methods, whether based on volume of milk analyzed or mass of 1 mL of milk analyzed. All values presented in the main text are in terms of ng/mL unless otherwise noted.

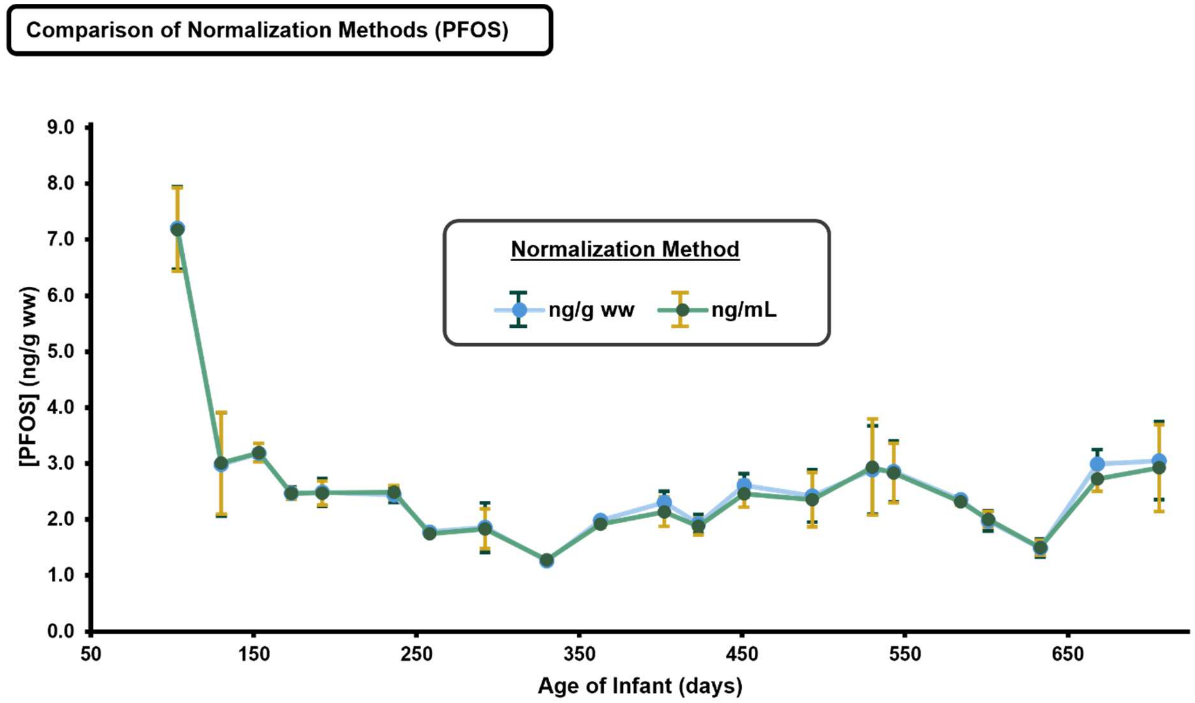

**Figure S6.** Analysis of batch-to-batch variability for pooled dolphin samples. Calculated concentrations of PFAS for each pooled sample (1 per sample preparation batch) determined to be above the LOQ for the analyte were used to calculate RSD across four biological replicates. The RSD (%) for these samples and analytes was less than 25%, where 30% was the predetermined threshold for evaluating batch-batch variability and is generally accepted to be the deviation attributable to biological sampling.

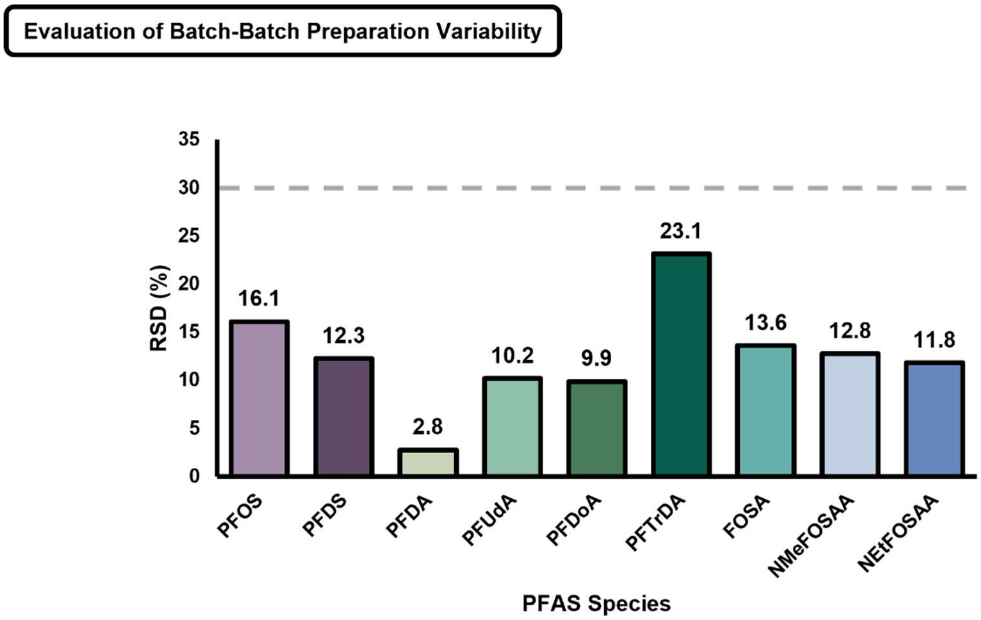

**Figure S7.** Determination of quantitation accuracy to NIST SRM 1954 (Fortified Human Breastmilk). This standard reference material does not list PFAS as a quantified class of compounds on its certificate of analysis, however a paper published by JM Keller *et al*. in the journal of Analytical Bioanalytical Chemistry (2010) published separate values obtained for a variety of PFAS, however values obtained by separate laboratories are incongruent with one another. For the two compounds (PFHpA and PFOS) with reported concentrations above the LOQs of this study, only PFOS has a "Consensus Value" across all laboratories, which is represented by the shaded purple bar, with error bars representing the "Expanded Uncertainty." For PFHpA, only one institution was able to quantify this analyte, whose average is represented by the shaded green bar, with error bars representing one standard deviation. Throughout method development, concentrations calculated for the NIST SRM were consistently lower than the reported value. These values match what is reported by authors of other studies (Abafe *et al. Molecules* (2021)). We acknowledge that this under-representation of the PFAS concentrations could be due to biological variability, or inadequate characterization of the SRM by previous studies.

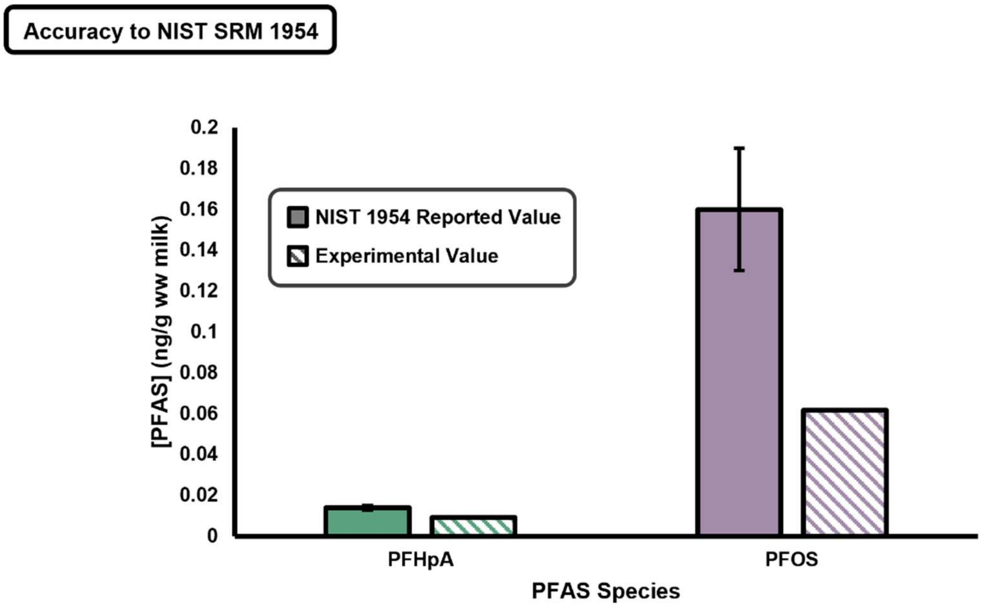

**Figure S8.** Definitions of LOQ and MDL for quantitative and non-targeted analyses.

| **For Quantified Analytes** | |
|---|---|
| **Limit of Quantitation (LOQ)** | • LOQ was defined as the lowest point on the calibration curve which has a maximum CV (RSD) of 20% |
| **Method Detection Limit (MDL)** | • This study follows recommendation from the EPA based on method blanks (EPA 821-R-16-006, 2016)<br>• MB is defined as the **calculated concentration** of PFAS species identified in goat milk samples spiked with internal standard and taken through the extraction process.<br>1. If none of the method blanks yield a peak area >0 for an individual analyte, MDL is defined as the lowest detectable concentration following data processing and demultiplexing.<br>2. If some (but not all) of the method blanks for an individual analyte give peak areas >0, the MDL will be defined as the highest method blank (MB) concentration.<br>3. If detected in all of the method blanks, calculate as the $X_{MB}$ + 3*$SD_{MB}$, where X is the average concentration of the method blanks. |

| **For All Other Analytes**<br>(Targeted, Suspect Screening and Trendline Additions) | |
|---|---|
| **Method Detection Limit (MDL)** | • This study follows recommendation from the EPA based on method blanks (EPA 821-R-16-006, 2016)<br>• MB is defined as the **peak area** of PFAS species identified in goat milk samples spiked with internal standard and taken through the extraction process.<br>1. If none of the method blanks yield a peak area >0 for an individual analyte, MDL is defined as the lowest detectable peak area following data processing and demultiplexing.<br>2. If some (but not all) of the method blanks for an individual analyte give peak areas >0, the MDL will be defined as the highest method blank (MB) peak area.<br>3. If detected in all of the method blanks, calculate as the $X_{MB}$ + 3*$SD_{MB}$, where X is the average peak area of the method blanks. |

**Figure S9.** Quantified PFAS over lactation course separated by sampling date.

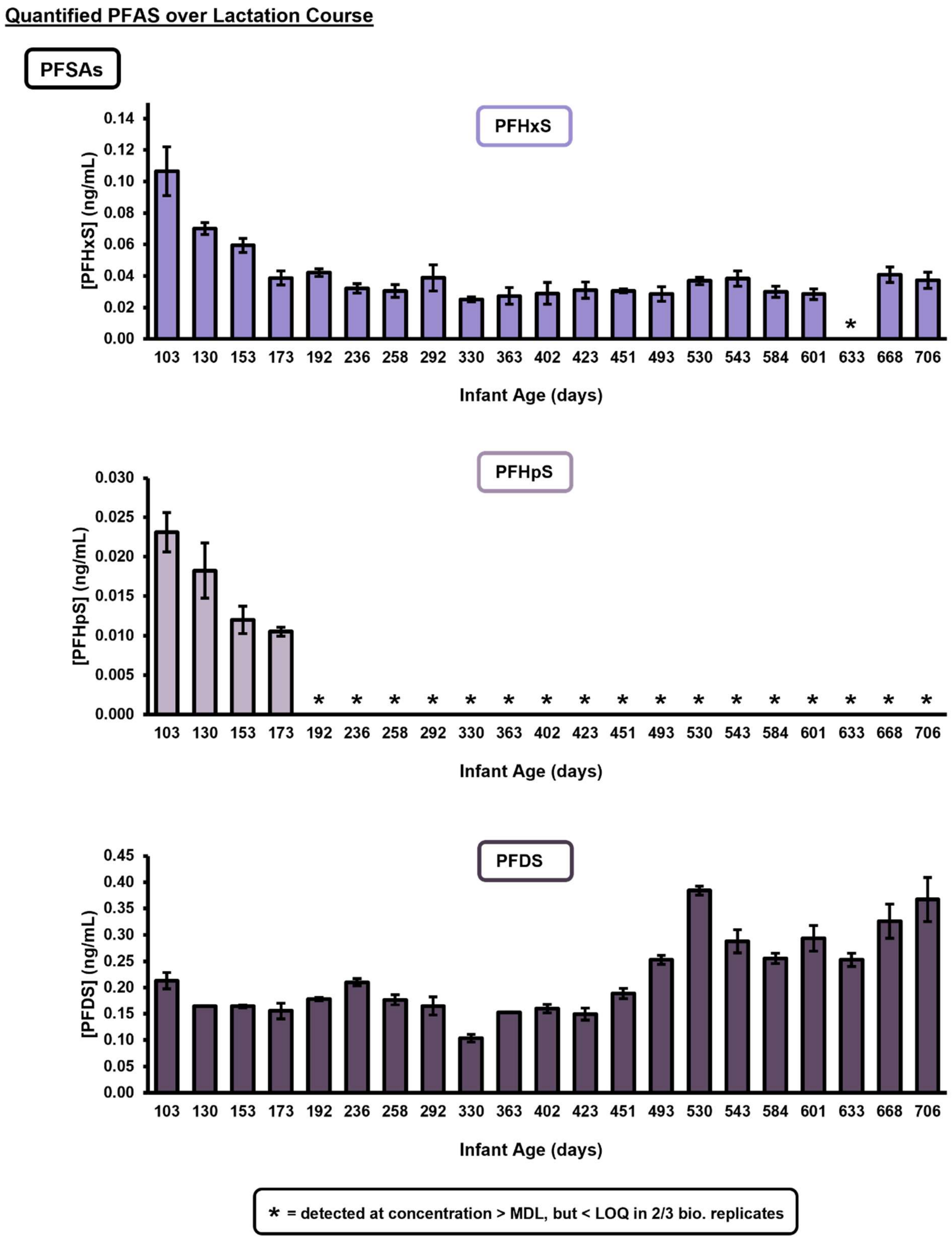

**PFCAs**

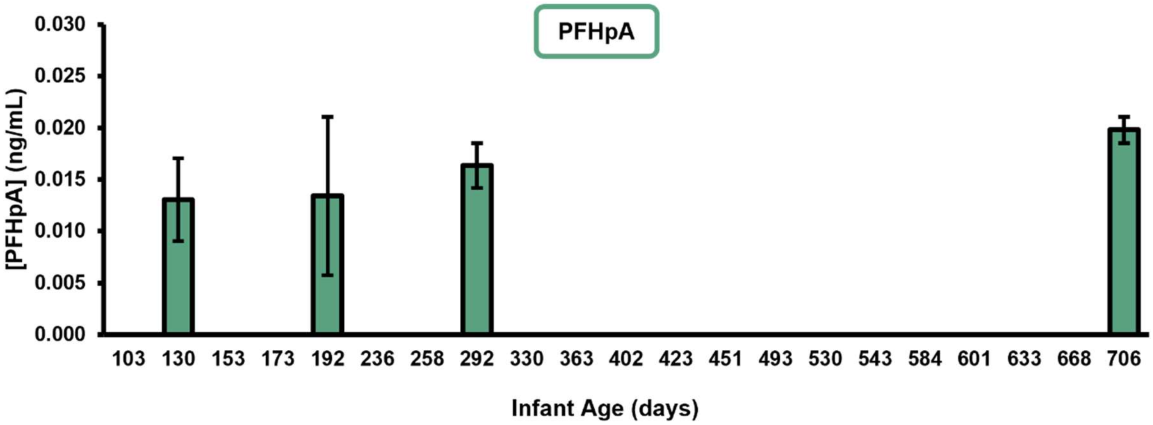
PFHpA
[PFHpA] (ng/mL)
0.030
0.025
0.020
0.015
0.010
0.005
0.000
103 130 153 173 192 236 258 292 330 363 402 423 451 493 530 543 584 601 633 668 706
Infant Age (days)

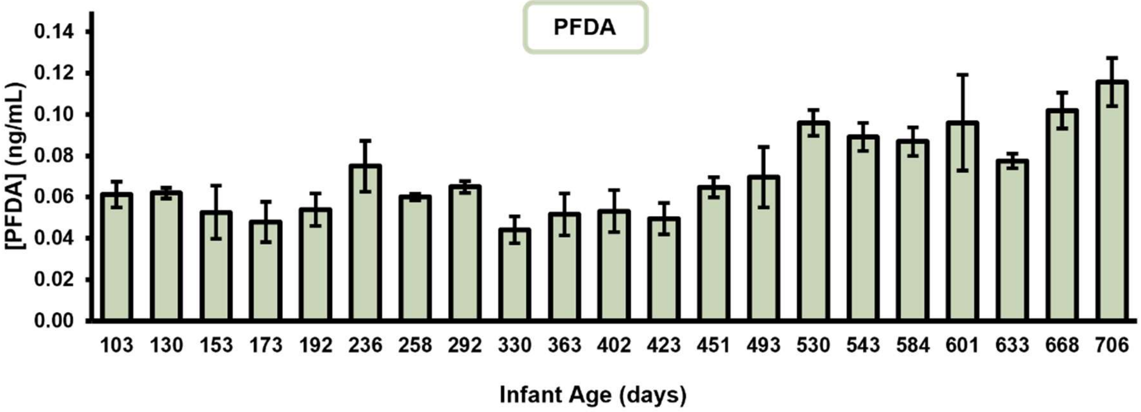
PFDA
[PFDA] (ng/mL)
0.14
0.12
0.10
0.08
0.06
0.04
0.02
0.00
103 130 153 173 192 236 258 292 330 363 402 423 451 493 530 543 584 601 633 668 706
Infant Age (days)

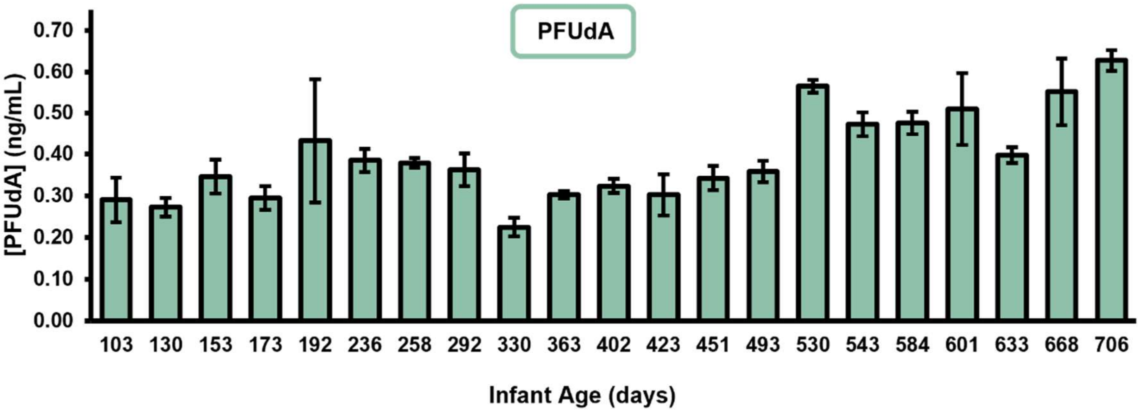
PFUdA
[PFUdA] (ng/mL)
0.70
0.60
0.50
0.40
0.30
0.20
0.10
0.00
103 130 153 173 192 236 258 292 330 363 402 423 451 493 530 543 584 601 633 668 706
Infant Age (days)

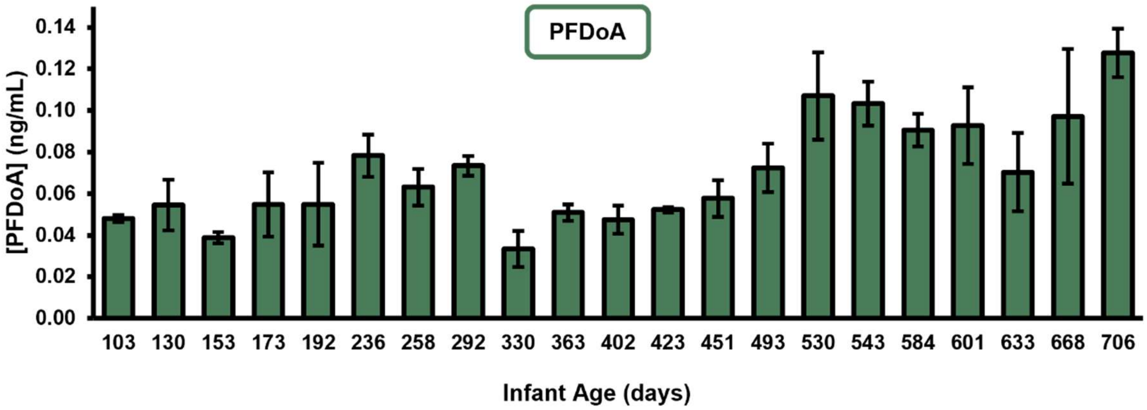


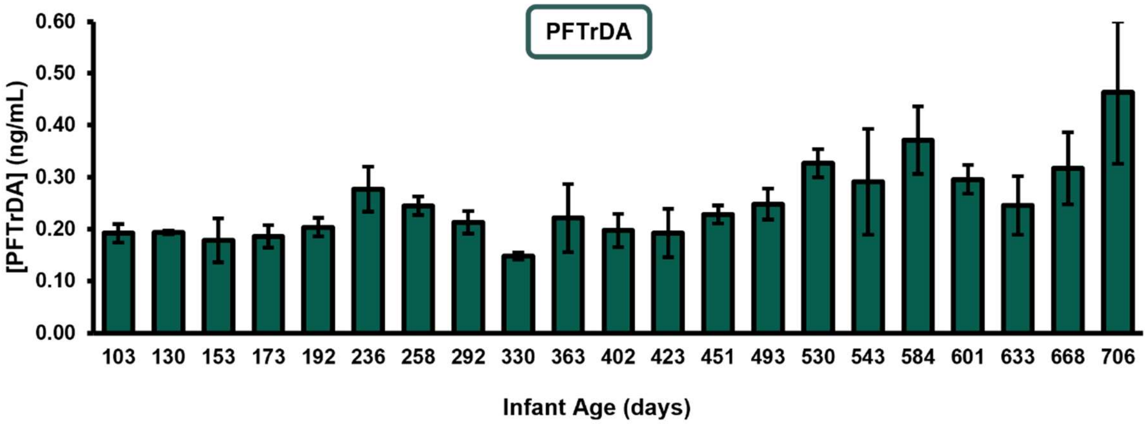


**FTSs**

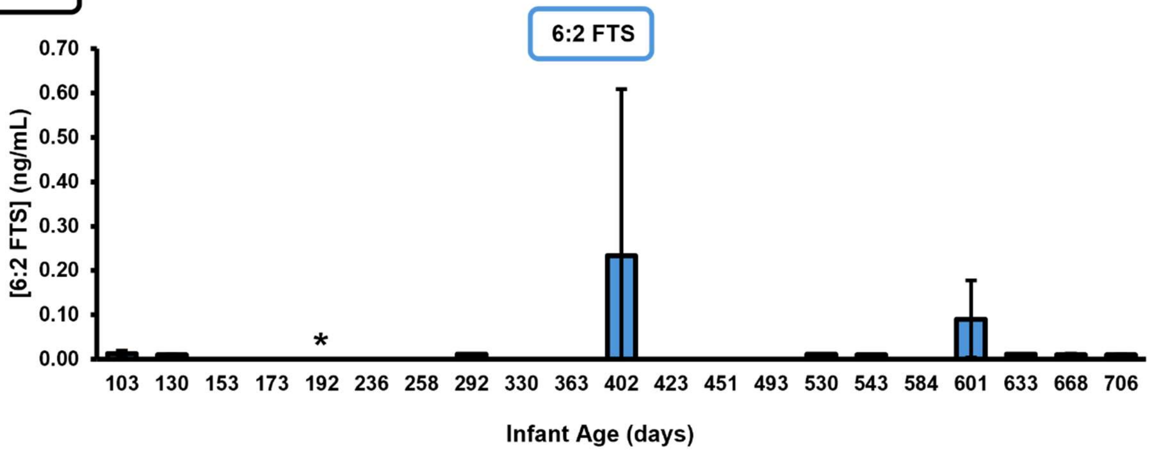


* = detected at concentration > MDL, but < LOQ in 2/3 bio. replicates

**PFASAs**

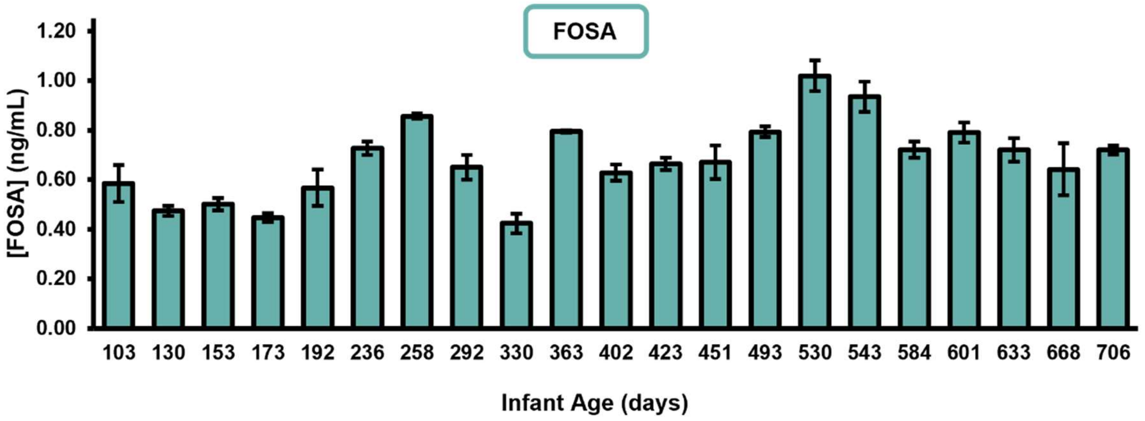


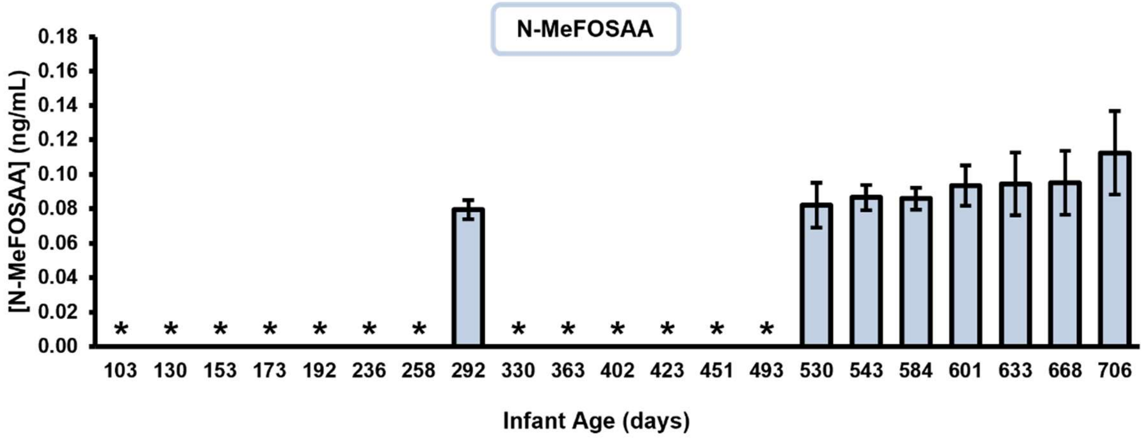


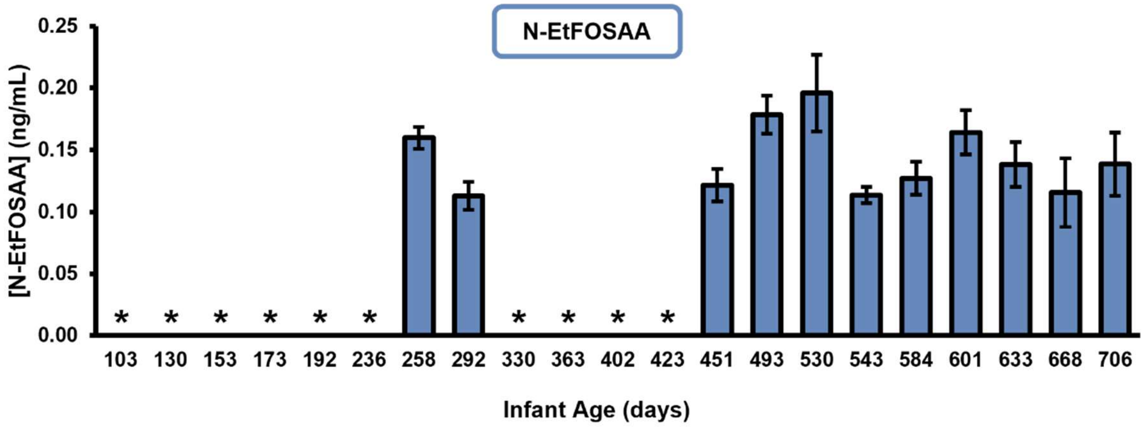


* = detected at concentration > MDL, but < LOQ in 2/3 bio. replicates

**Figure S10.** Example calculation for weekly PFAS intake at three months of age.

**Estimating PFOS Weekly Intake at 3 Months**

From Flower *et al.* Table 5. At Month 3, neonate was fed 285-590 mL of formula with a feed frequency of "Weaned from [every 2 hours] to [every 3 hours]. During this month, the dolphin weighed between 32.4-47.0 kg. This study found that at 3 months of age, (approx. sampling date 103, the mean PFOS concentration in milk would be 7.18 ng/mL.

$$\text{Daily Formula Intake} = \frac{1\ Feeding}{Average\ \ 2.5\ hours} \times \frac{24\ hours}{1\ day} \times \frac{Average\ 437.5\ mL}{1\ Feeding} = 4{,}200\ \frac{mL}{day}$$

$$\text{Weekly Formula Intake} = \frac{4{,}200\ mL\ formula}{day} \times \frac{7\ days}{1\ week} = 29{,}400\ \frac{mL}{week}$$

$$\text{Weekly PFOS Intake} = \frac{29{,}400\ mL}{week} \times \frac{7.18\ ng\ PFOS}{mL\ milk} \times \frac{}{Average\ 39.7\ kg\ bw} \approx 5316\ \text{ng PFOS/kg bw/week}$$

**Figure S11.** Percentages of branched isomer for quantified PFAS over course of lactation.

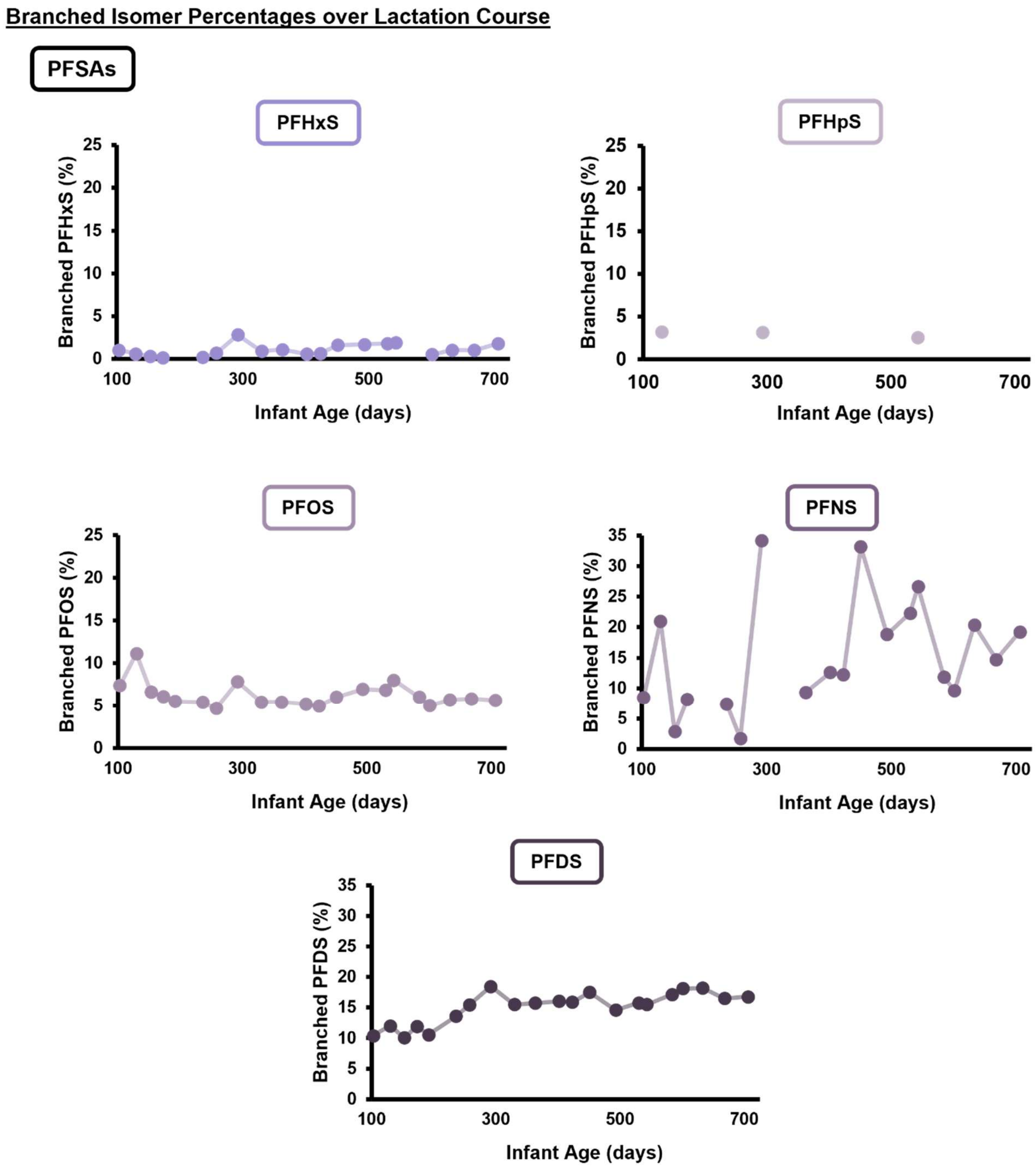

#### Branched Isomer Percentages over Lactation Course cont.

**PFCAs**

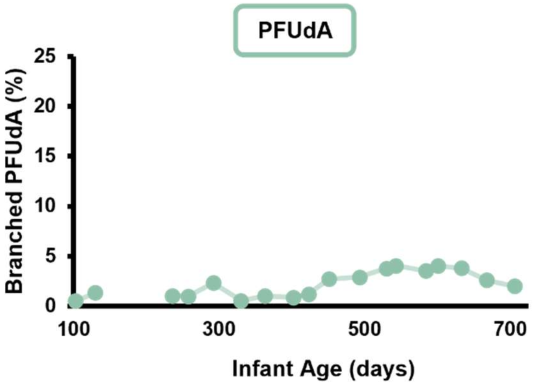


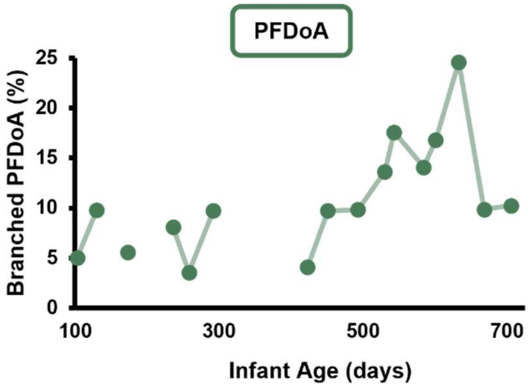


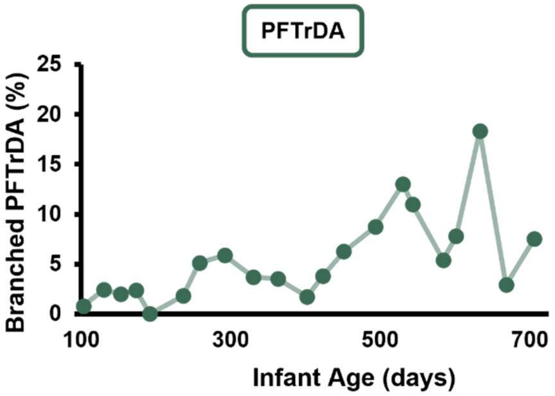


**PFASAs**

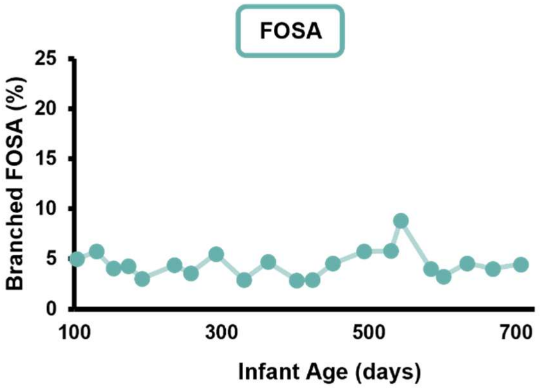

**Figure S12.** Two long-chain PFSAs (perfluoroundecane sulfonic acid (PFUnS) and perfluorotridecane sulfonic acid (PFTrS)) were identified using CCS vs. *m/z* and RT vs. *m/z* trendlines. Trendlines were drawn based on CCS and RT values from library based on standards (purple points). CCS and RT values for PFUnS and PFTrS are experimentally determined from a replicate injection from individual sampling dates.

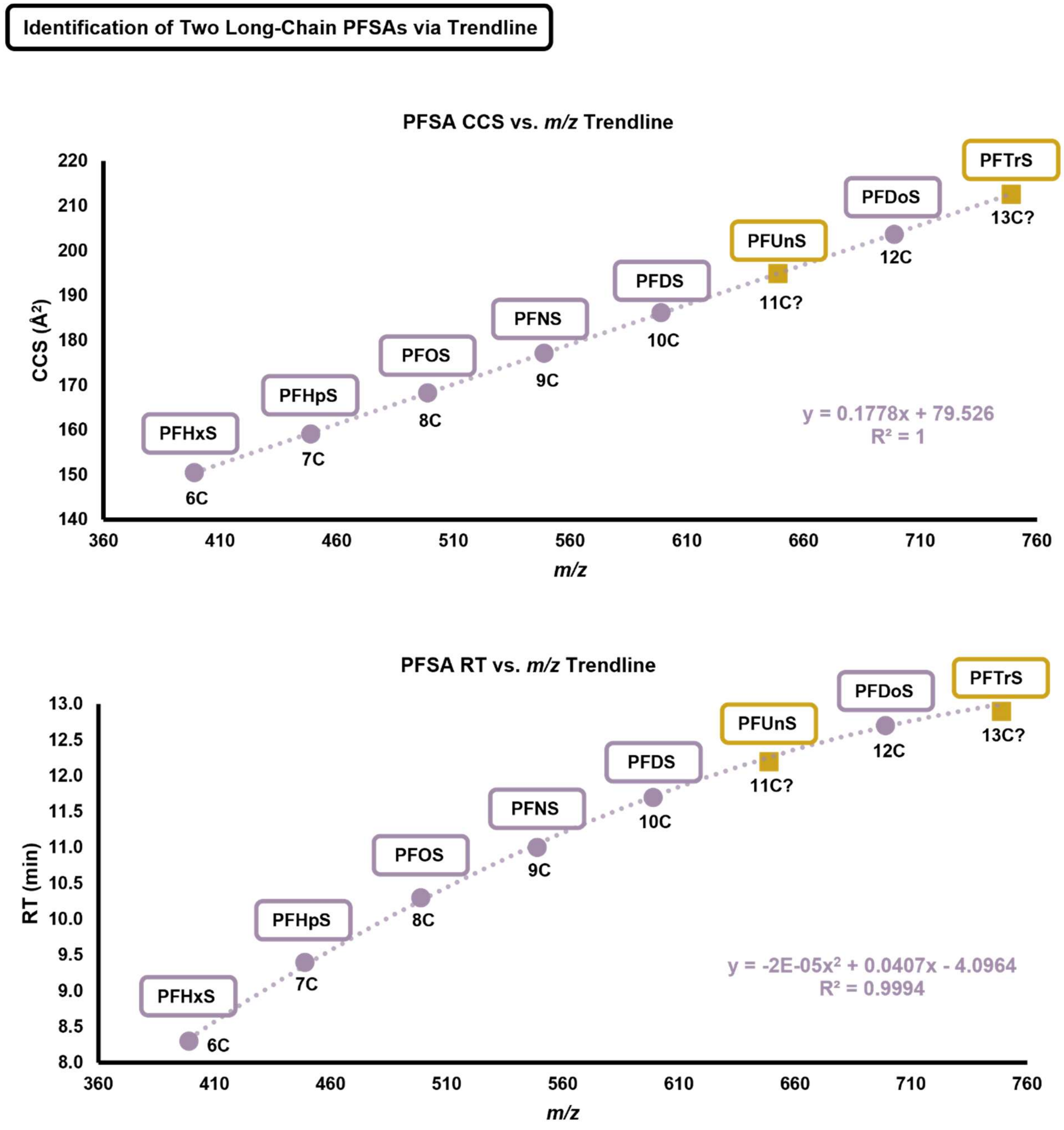
